# Polo-like kinase 1 controls activation of adult neural stem cells and represents a druggable entry-point to boost adult neurogenesis

**DOI:** 10.64898/2026.05.12.724486

**Authors:** Ana L. Barrios-Muñoz, Coral López-Fonseca, Elena Gutiérrez-Galindo, Jose Manuel Morante-Redolat, Antonio Jordán-Pla, Btissam Ben-Said, María Arroyo-Camuñas, Alonso M. Higuero, Francisco Zafra, Teresa Iglesias, Isabel Fariñas, Marcos Malumbres, Eva Porlan

**Affiliations:** Departamento de Biología Molecular and Instituto Universitario de Biología Molecular (IUBM), Universidad Autónoma de Madrid. Madrid, Spain; Centro de Biología Molecular Severo Ochoa (CBM), Consejo Superior de Investigaciones Científicas (CSIC)-UAM, Madrid, Spain. Madrid, Spain; Centro de Investigación Biomédica en Red de Enfermedades Neurodegenerativas (CIBERNED), Instituto de Salud Carlos III. Madrid, Spain; Instituto de Biotecnología y Biomedicina (BioTecMed), and Departamento de Biología Celular y Biología Funcional. Universidad de Valencia. Valencia, Spain; Membrane Biology and Axonal Repair Laboratory. Hospital Nacional de Parapléjicos (SESCAM) and Instituto de Investigación Sanitaria de Castilla-La Mancha (IDISCAM). Toledo, Spain; Instituto de Investigación Sanitaria del Hospital Universitario La Paz (IdiPAZ), Instituto de Salud Carlos III. Madrid, Spain; Instituto de Investigaciones Biomédicas Sols-Morreale (IIBM) CSIC-UAM, Madrid, Spain; Vall d’Hebron Institute of Oncology (VHIO). Barcelona, Spain; Institució Catalana de Recerca i Estudis Avançats (ICREA). Barcelona, Spain; Centro de Investigación Biomédica en Red de Cáncer (CIBERONC), Instituto de Salud Carlos III. Madrid, Spain

**Author notes:** **Address for correspondence:** Eva Porlan. Equal contribution (Ana L. Barrios-Muñoz, Coral López-Fonseca). AstraZeneca Farmacéutica S.A. Spain. Spanish National Cancer Research Centre (CNIO), Madrid, Spain.

## Abstract

Neural stem cells (NSCs) persist in the mammalian brain after development, but their potential for replacing neural populations lost in brain damage is dramatically restricted. Quiescence, or the ability to transition in and out of the cell cycle is a hallmark of adult NSCs. It deepens as the organism ages and is regarded as a strategy to protect the reservoirs of NSC for the long term. Deciphering the molecular code of dormancy is a major interest in cellular and molecular biology of ageing and in regenerative medicine since it has important implications for potentially manipulating NSCs to improve tissular homeostasis. In this work we have identified the mitotic kinase Polo like kinase 1 (Plk1) as a novel intrinsic regulator of the dynamic transition between NSC quiescence and activation in adult neurogenic niches. Using quantitative phosphoproteomics to pinpoint mechanistical mediators of the role of Plk1, we have identified the quiescence/activation molecular switch formed by E3 ubiquitin ligase Huwe1 and its target, the transcription factor Ascl1/Mash1, as novel effectors of Plk1 in NSCs. We discover that Plk1 restrains the accumulation of Ascl1/Mash1 by acting upstream of Huwe1, providing a direct molecular connection between completion of the cell cycle and the re-establishment of a low activation state following NSC division, which supports the concept that quiescence is an actively maintained state. Moreover we demonstrate that pharmacological inhibition of Plk1 is sufficient to enhance adult neurogenesis in vivo, identifying Plk1 as a viable target to acutely boost endogenous stem cell activation.

## INTRODUCTION

The post-developmental generation of neurons is sustained by a pool of embryonic radial glial (RG) cell-derived adult neural stem cells (NSCs) that persist in niches within the mammalian brain (Bond et al., 2015), namely the subependymal zone (SEZ) of the lateral ventricle wall, and the subgranular zone (SGZ) of the dentate gyrus (DG). NSCs also persist in the human brain and can respond to certain injuries, making them a potential therapeutic target in regenerative medicine and ageing However, they appear to become largely inactive after infancy, which greatly restricts their capacity to replace neural populations lost following brain damage (Doludda et al., 2026).

In a persistently active niche such as the murine SEZ, NSCs give rise to progenitor cells through strictly restrained cell divisions. Progenitors in turn, divide rapidly to amplify the population and produce neuroblasts that migrate into the olfactory bulbs (OB), where they cycling at the time of labelling and differentiate into several types of interneurons (Lim and Alvarez-Buylla, 2016). Adult and developmental neurogenesis share mechanisms that range from the molecular control over cell cycle, cell-fate acquisition, survival, differentiation migration, maturation and the final integration of newborn neurons. However, one important difference between embryonic RG and NSCs is the latter’s need to regulate quiescence. The quiescent state (G0) is not simply a passive default in response to lack of mitogenic signals; rather, it is a reversible tightly controlled and still not fully understood condition (de Morree and Rando, 2023). In the mouse SEZ, adult NSCs can be found anywhere along a spectrum of quiescence, ranging from deep dormancy to a poised state ready to enter the cell cycle, and once activated, they can transition back out of the cell cycle (Basak et al., 2018; Belenguer et al., 2021). The reversible cell cycle arrest shields long-lived cellular populations from premature depletion and accumulation of DNA damage that could promote disruption of tissue homeostasis and/or oncogenic transformation, increasing as the organism ages (de Morree and Rando, 2023; Rando et al., 2025).

The molecular cues controlling the dynamic transition into and out of the cell cycle remain poorly characterized. The pro-activation basic helix–loop–helix transcription factor (bHLH) Achaete-scute homolog 1/mammalian achaete-scute homolog 1 (Ascl1/Mash1) and its proteolytic regulation by the HECT, UBA and WWE domain containing E3 ubiquitin protein ligase 1 (Huwe1) is one of the few described mechanisms ruling the transit between quiescence and activation (Urbán et al., 2016), but it has not been linked to cell cycle control so far. The Ser/Thr kinase Polo like kinase (Plk1) is a master regulator of cell division. Through its activity on several substrates, Plk1 controls centrosome maturation, assembly of the spindle and mitotic entry and exit, chromosome segregation and cytokinesis (Barr et al., 2004). It also controls the proper mode of division of RG during development (Connell et al., 2017; Gonzalez-Martinez et al., 2022; Sakai et al., 2012; Wachowicz et al., 2016). Its role in adult NSCs, which in contrast to RG, proliferate mainly symmetrically and need to additionally balance activation with their prevalent state of quiescence (Obernier et al., 2018), has not been studied.

In this work we have found that Plk1 activity prevents the accumulation of Ascl1/Mash1 in NSCs by acting upstream of its proteostatic regulator Huwe1. Our findings indicate that in mitosis Plk1 acts as a mechanistic link connecting Ascl1/Mash1 degradation to the end of the cell cycle, ensuring the restoration of pro-activation protein levels following cell division. Through this mechanism, Plk1 regulates the likeliness of NSC re-activation and serves to restrain the post-developmental neuronal output, which in turn can be enhanced artificially by means of small molecule inhibitors of the kinase. We identify a previously undescribed functional interaction between Plk1 and Ascl1/Mash1 via Huwe1 in both neurogenic niches, and pinpoint Plk1 as a key mechanistical player in the timely activation of NSCs. Additionally, we uncover a novel potential of NSCs to be pharmacologically regulated on demand.

## RESULTS

### Plk1 restrains entry of adult quiescent NSCs into activation

To investigate Plk1 functions in neurogenesis, we started by assessing *Plk1* expression in the different populations of the SEZ neurogenic lineage. We made use of a previously published public RNA-seq dataset (GEO: GSE138243) (Belenguer *et al*., 2021) obtained by the stratification and isolation by fluorescence-activated cell sorting (FACS) of the SEZ lineage populations *as per* the strategy outlined (Fig. S1A). In this study, non-relevant populations (oligodendrocytes, erythrocytes, microglia and endothelial cells) were filtered out by positivity to a combination of markers. Lineage negative cells were then immunostained with a set of markers for SEZ cells (Fig. S1B) allowing for the transcriptomic characterization of NSCs in different states of activation (quiescent (q), primed (p) and active (a) NSCs), together with neural progenitor cells (NPCs) and neuroblasts (NBs) (Belenguer *et al*., 2021). Not unexpectedly for a cell cycle regulator, *Plk1* expression correlated mainly with proliferating populations, being increased in aNSCs, highly proliferative NPCs 1 and 2, and in the proliferating early NB subgroup (NB1), before it decreased in the postmitotic population of migrating late NB2 (Fig. 1A). Interestingly, in the quiescent NSC populations, its levels were relatively low but started to increase in pNSCs. Aside from its potential effects in NSC cycling, the gradual increase of *Plk1* expression across the quiescence-to-activation transition prompted us to hypothesize that Plk1 could be involved in the activation of NSCs from quiescence.

**Figure 1.**
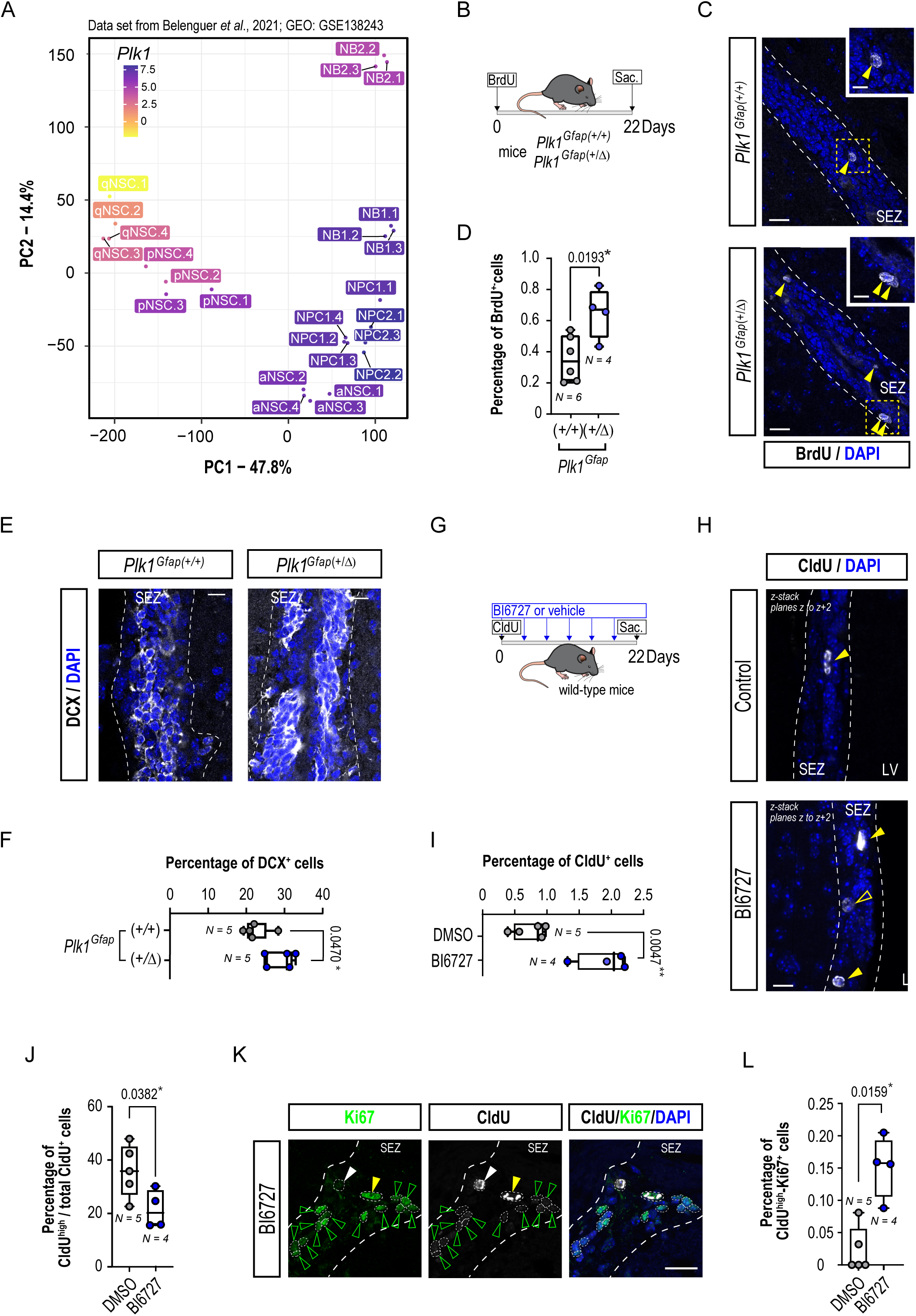
Plk1 restrains entry of adult quiescent NSCs into activation. **(A)** Expression of Plk1 in SEZ populations. Principal component analysis (PCA) of bulk RNA-seq profiles from SEZ cell populations and their biological replicates (Belenguer *et al*., 2021). The first two components account for 62.2% of the total variance, with samples and labels colored by log-normalized *Plk1* expression. qNSC, quiescent NSC; pNSC, primed-to-activate NSC; aNSC, active NSC; NPC1 and 2, neural progenitor cell populations 1 and 2; NB1 and 2, neuroblast population 1 and 2. Dataset GEO: GSE138243**. (B)** Schematic depicting the BrdU injection-chase protocol used in *Plk1^Gfap(+/+)^* and *Plk1^Gfap(+/^Δ)* mice. **(C)** Confocal micrographs of the staining for BrdU^+^-retaining cells (gray, yellow arrowheads) in the SEZ (delimited by white dashed lines) of *Plk1^Gfap(+/+)^*and *Plk1^Gfap(+/^Δ)* mice. Nuclei are stained with DAPI (blue). Scales: 20 μm and 10 μm (insets). (**D**) Quantification of the percentage of BrdU^+^ cells in the SEZ of *Plk1^Gfap(+/+)^*(N = 6) and *Plk1^Gfap(+/^Δ)* (N = 4) relative to the total number of cells. (**E**) Confocal micrographs of the SEZ *Plk1^Gfap(+/+)^* and *Plk1^Gfap(+/Δ)^* mice, immunostained for the neuroblast marker DCX (gray). Nuclei are stained with DAPI (blue). Scale bars: 20 μm. Dashed white lines delimit the boundaries of the SEZ. (**F**) Quantification of the percentage of DCX^+^ cells in the SEZ of *Plk1^Gfap(+/+)^* (N = 5) and *Plk1^Gfap(+/^Δ)* (N = 5) mice. **(G)** Schematic depicting the CldU and BI 6727 injection-chase protocol used in wild-type mice. (**H**) Confocal micrographs of the SEZ (white dashed lines) of control or BI 6727-treated mice immunostained for CldU (gray, yellow arrowheads). Full yellow arrowheads point at CldU^high^ cells. Cell nuclei are stained with DAPI (blue). Scale bars: 20 μm. LV, lateral ventricle. (**I**) Quantification of the percentage of CldU^+^ cells in the SEZ of vehicle-(control, N = 5) or BI 6727-treated mice (N = 4). (**J**) Percentage of high intensity CldU^+^ (CldU^high^) within the total of CldU^+^ cells in the SEZ of vehicle- (control, N = 5) or BI 6727-treated mice (N = 4). (**K**) Example confocal micrographs of the co-staining for BrdU (gray) and the proliferation marker Ki67 (green) in a BI 6727-treated wild-type mouse. The white dashed lines delimit the SEZ. The white, full arrowhead points at a BrdU^+^-Ki67^-^ cell, the green arrowheads point at a BrdU^-^-Ki67^+^ cells, and one activated NSC is identified as BrdU^+^-Ki67^+^ double positive cell (yellow, full arrowhead). Scale bars: 20 μm. (**L**) Percentage of cycling (Ki67^+^)-high intensity CldU^+^ (CldU^high^) cells in the SEZ of vehicle- (control, N = 5) or BI 6727-treated mice (N = 4). Data are represented as box and whiskers from min. to max and all data points are shown. \**p*< 0.05 by two-tailed unpaired *Student*’s *t*-tests (**D, F, I, J**) and *Mann-Whitney*’s test (**L**). Symbols represent biological replicates (**A**) or mean data from individual mice (**D, F, I, J, L**).

Because of the essential role of Plk1 during mitosis, we used heterozygous mice to study the functional effects of the kinase in adult NSCs. The mice were generated by crossing *Plk1^(+/lox)^*(Wachowicz *et al*., 2016) mice with a strain expressing *Cre* downstream the murine *Gfap* promoter (Garcia et al., 2004) to conditionally delete *Plk1* in NSCs and derived progeny (*Plk1^Gfap^*^(+/Δ)^) (Fig. S2A). *Plk1^Gfap^*^(+/Δ)^ and control *Plk1^Gfap^*^(+/+)^ mice were repeatedly administered the thymidine analogue bromodeoxyuridine (BrdU) on the same day and sacrificed after a 3-week chase (Del Puerto et al., 2023) (Fig, 1B). This paradigm mostly serves to assess the numbers of NSCs that were cycling at the time of labelling and did not dilute the nucleoside during the chase by re-acquiring quiescence. While *Plk1^Gfap^*^(+/Δ)^ mice presented a two-fold increase in the percentage of BrdU^+^-retaining cells *vs*. control mice (Fig. 1C, D), we also found an evident increase in NB production, determined as DCX^+^ cells (Fig. 1E, F).

To address this apparent paradox, we decided to modify Plk1 activity in a time-controlled manner by using a specific small-molecule inhibitor (iPlk1, BI 6727) (Danovi et al., 2013; Rudolph et al., 2009). We first labeled a cohort of cycling cells in wild-type mice with chlorodeoxyuridine (CldU) and then treated them with BI 6727 or vehicle every three days for 3 consecutive weeks (Fig. 1G). Since fully inhibiting Plk1 halts cell division in neural progenitors (Gonzalez-Martinez *et al*., 2022), we used a low dose of BI 6727, and we made sure that the treatment did not provoke a mitotic arrest in the SEZ by immunostaining for the cell cycle marker Ki67 (Fig. S2B, C). Also, in line with our data from the genetic approach, we observed that the pharmacological inhibition of Plk1 provoked a two-fold increase in the percentage of cells retaining CldU in the SEZ (Fig. 1H, I). However, when we scored the proportions of cells with the highest undiluted CldU label (CldU^high^), over all CldU^+^ cells, we found that BI 6727-treated mice displayed fewer of these cells (Fig. 1J). This suggested that a higher proportion of CldU-retaining cells might have been re-entering proliferation in mice with lower Plk1 activity. Accordingly, in BI 6727-treated mice, more CldU^+^ cells were also positive for the proliferation marker Ki67 (Fig. 1K, L). Together, our results indicated that quiescent NSCs had a higher probability to re-engage in cell cycle when Plk1 was inhibited. These data reconciled with the results obtained in the genetic model and indicated that Plk1 restrains activation of NSCs.

Our *in vivo* results prompted us to test whether Plk1 inhibition could indeed facilitate the entry of NSCs into activation by using neurosphere cultures. When isolated from the SEZ and plated *in vitro*, qNSCs do not survive whereas both pNSCs and aNSCs contribute to the formation of neurospheres, although at different paces (Belenguer *et al*., 2021). We generated primary neurosphere cultures from *Plk1^(+/lox)^* mice crossed with *Pol2a-Cre*ERT2 strain that expresses Cre ubiquitously (Wachowicz *et al*., 2016), herein called *Plk1^(+/lox)^* mice. Treatment of freshly dissociated SEZ cells with 4-OH-tamoxifen (Tam) to acutely reduce Plk1 provoked an increase in the number of primary neurospheres generated (Fig. 2A). In contrast, the effect of overexpressing *Plk1* was the opposite: generation of primary neurospheres from mice with a knock-in allele [tet_*PLK1*(T)] (de Carcer et al., 2018) in which a FLAG-tagged version of hPLK1 can be induced upon the addition of doxycycline (Dox) (Fig. S3A), herein called Plk1*^(+/Ki,)^*, was severely decreased in Dox-treated cultures *vs*. vehicle (Fig. S3B). Combined, our data indicated that Plk1 levels determine the likelihood of an NSC to reach the activation entry point.

**Figure 2.**
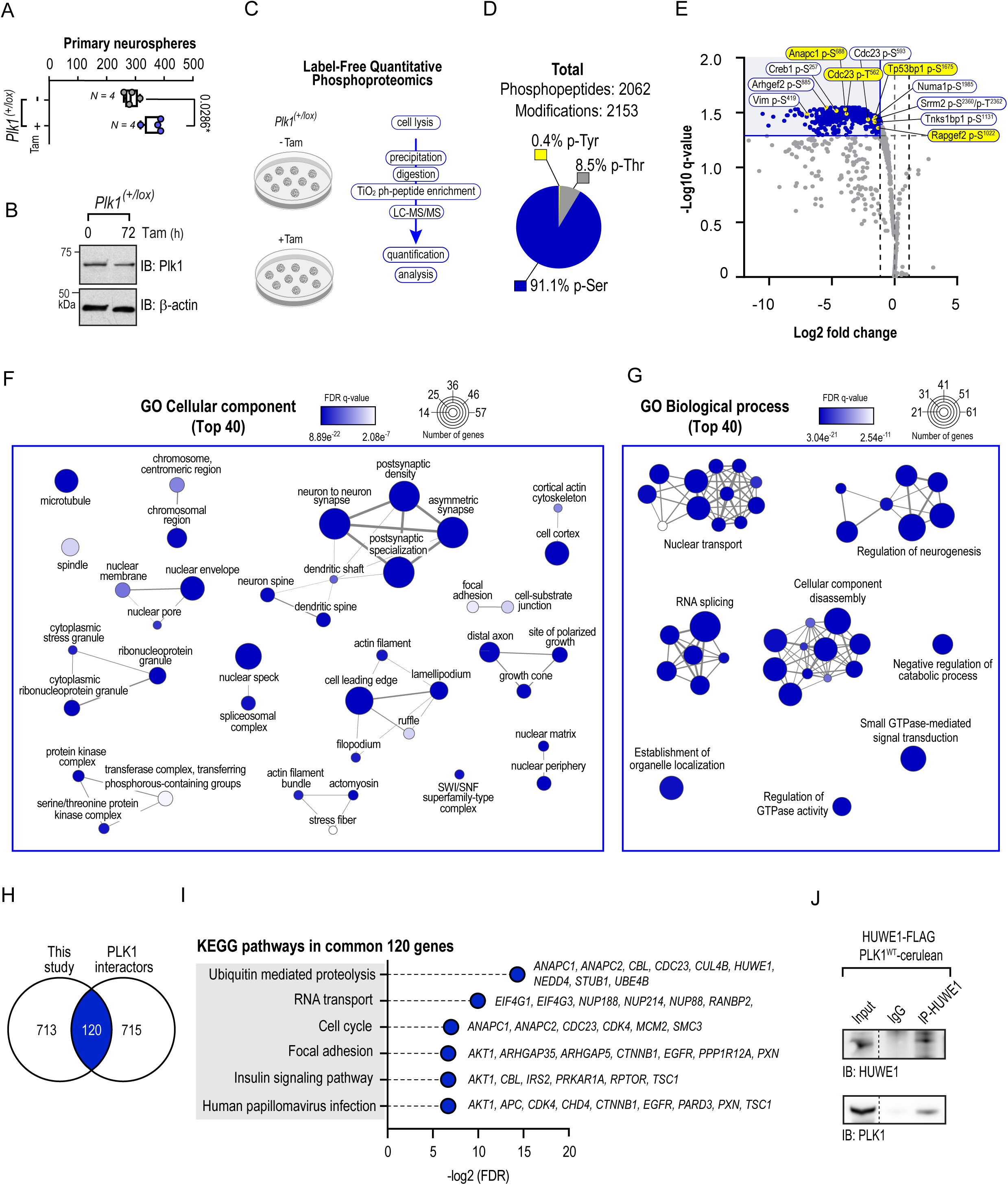
Unbiased phosphoproteomics identifies a link between Plk1 and HUWE1 in NSCs. **(A)** Quantification of the number of neurospheres obtained from freshly disaggregated SEZ tissue from *Plk1^(+/lox)^* mice, treated with vehicle- (N = 4) and 4-OH-tamoxifen (Tam) (N = 4). Data are represented as box and whiskers from min. to max and all data points are shown. \**p* < 0.05, by *Mann-Whitney*’s test. Dots represent independent cell cultures. (**B**) *Western* blot for Plk1 and β-actin as loading control in lysates from control or Tam-treated neurospheres from *Plk1^(+/lox)^* mice. **(C)** Schematic depicting the pipeline followed in the label-free differential analysis of phosphoproteins by liquid chromatography coupled with tandem mass spectrometry performed in *Plk1^(+/lox)^* vehicle-(N = 4) and Tam-treated (N = 4) neurospheres (**D**) Identified phosphorylation distribution by residue type. (**E**) Volcano plot showing all phosphopeptides identified in ≥ 2 replicates. Blue lines and background indicate the sector of the plot that include the cut-off values (*q* value = 0.05, and Log2 _[Tam/Vehicle]_ = ± 1.0). Yellow dots indicate known Plk1-dependent (indicated in yellow boxes) or Plk1-inhibitor-sensitive phosphosites (in white boxes). In every case the position of the residue corresponds to the mouse protein. (**F, G**) Over-representation analysis network representation of the 40 most differentially regulated genes found in Tam-treated NSCs *vs*. control. Gene ontologies (GO)-cellular component (**F**) and biological processes (**G**) are shown. Similar gene sets are grouped into functional modules. The node size represents the number of genes, the node colour correlates with the FDR q-value, and the edge width with the number of overlapping genes. (**H**) Venn diagram showing the genes from this study common to a list of PLK1 interactors (BioGRID). (**I**) KEGG pathway analysis of the list of 120 common genes, showing ‘Ubiquitin mediated proteolysis’ as the top pathway represented, and the gene names corresponding to each category. (**J**) Immunoprecipitation (IP) of HUWE1 showing co-IP with PLK1 in representative immunoblots (IB) for the specific detection of HUWE1 and PLK1 in lysates from transfected HEK293T cells. IgG, isotype control antibody. One experiment is shown out of three performed with similar results. Irrelevant lanes were cropped out of the image (dashed black lines). For full *Western* blot images refer to Fig. S7.

### Unbiased phosphoproteomics identifies a link between Plk1 and Huwe1 in NSCs

Since our functional data indicated that Plk1 appeared to restrain NSC exit from quiescence, we next set out to discover possible accountable mechanistical targets of Plk1 by performing a differential analysis of unlabelled phosphoproteins by liquid chromatography coupled with tandem mass spectrometry (LC-MS/MS). To capture potentially direct targets of Plk1 in adult SEZ NSCs we used neurosphere cultures from *Plk1^(+/lox)^*, treated with 4-OH-tamoxifen (Tam) or vehicle for 72h, that showed reduced Plk1 levels (Fig. 2B, C). The complete data set of the comparative phosphoproteomic analysis is provided in https://doi.org/10.5281/zenodo.20036962 and in Supplemental Spreadsheet 1. The events that corresponded to phosphopeptides and that were detected in ≥ 2 replicates summed 2,062, and 2,153 identified phosphoresidues, of which 91.1% were serines, 8.4% threonines and 0.4% tyrosines (Fig. 2D). After the quantitative analysis we identified 1,532 phosphopeptides with negative differential regulation *vs*. none with positive differential regulation. Among the negative differentially regulated phosphopeptides, we found several corresponding to previously known Plk1 targets and, as expected, we were able to detect known Plk1-dependent and Plk1-inhibitor-dependent phosphosites in these proteins (Fig. 2E). Next, we obtained a list of 833 unique proteins after using a Log2 [Plk1^(+/Δ)^/Plk1^(+/lox)^] < -1.0 cut-off (Supplemental Spreadsheet 2). Comparing this list to previously published PLK1-dependent phosphoproteomics analyses (Kettenbach et al., 2011; Oppermann et al., 2012), we found 50 common proteins to the 3 studies (Fig. S4A, B) further validating our assay. Then, we performed over-representation analysis using the full list of 833 gene-names. The enrichment map and network representation of the analysis for the top 40-GO terms related to cellular component (Fig. 2F) rendered several clustered terms, many linked to the canonical Plk1-dependent well-known control of mitosis and cytokinesis. However, in our analysis, the biggest cluster was related to synapse components, grouping genes such as microtubule associated proteins or actin-microtubule crosslinkers (Supplemental Spreadsheet 3). The map and network representation of the 40 biological processes with more differentially regulated gene-names found in Plk1-deficient NSCs rendered four clusters of terms, and four other unclustered terms (Fig. 2G). One of the biggest cluster was related to ‘regulation of neurogenesis’. Altogether the data indicated that some of the canonical functions of Plk1 were still represented in our analysis in NSCs cells, while other less known or previously undescribed biological processes in which the kinase might take part were likely present as well. At this point we decided to narrow down the list of our putative hits by comparing the full list of 833 proteins with a list of 835 PLK1 interactors from the open repository of protein interactions BioGRID (Stark et al., 2006). We obtained 120 proteins common to both lists (Fig. 2H, Supplemental Spreadsheet 4), on which we performed Kyoto Encyclopedia of Genes and Genomes (KEGG)-pathway enrichment analysis, and as the Top 1 pathway we obtained ‘Ubiquitin mediated proteolysis’ (Fig. 2I). Amongst the genes in this category, HECT, UBA and WWE domain containing E3 ubiquitin protein ligase 1 (Huwe1) has been described as a negative regulator of the NSC pro-activation protein and master regulator of neurogenesis, Ascl1 (Urbán *et al*., 2016). To confirm that Huwe1 interacts with Plk1, we performed co-immunoprecipitation experiments in transfected HEK293T cells supporting the physical association between both proteins (Fig. 2J). The specific Huwe1 phosphoresidues found differentially regulated in our analysis are listed in Fig. S4C.

### Plk1 functions upstream of the Huwe1-dependent regulation of Ascl1 during mitosis

Interestingly, *in vitro* Ascl1 could be increased by the pharmacological inhibition of Plk1 (Fig. 3A, B) and by genetically silencing Plk1 (Fig. 3C left panel, D upper panel), or on the contrary, decreased upon Plk1 overexpression (Fig. 3C right panel, D bottom panel), in lysates from *Plk1^(+/lox)^* and *Plk1^(+/KI)^* treated with Tam or Dox, respectively. Cyclins D1 and D2 are targets of Ascl1 in NSCs (Urbán *et al*., 2016), so we evaluated whether the levels of the transcription factor correlated directly with its targets’ abundance in our Plk1 loss- and gain-of-function models. We found increased levels of cyclin D2 in *Plk1^(+/lox)^*NSCs treated with Tam, in contrast to *Plk1^(+/KI)^* NSCs treated with Dox, which displayed a decrease in cyclin D2 levels, both models showing a direct correlation with Ascl1, and inverse with Plk1 (Fig. 3C, E). Next, in HEK293T cells transfected with *cerulean* (*cer*)-tagged human *PLK1* versions (wild-type PLK1, kinase-dead PLK1^K82R^ and constitutively active PLK1^T210D^) together with *myc*-tagged *Ascl1*, we found that the Plk1-induced reduction of Ascl1 levels was independent from the latter’s transcription and dependent on the former’s catalytic activity, since constitutively active PLK1^T210D^-cer provoked a significant decrease of Ascl1-*myc* compared with wild-type and kinase-dead versions of the kinase (Fig. 3F, G).

**Figure 3.**
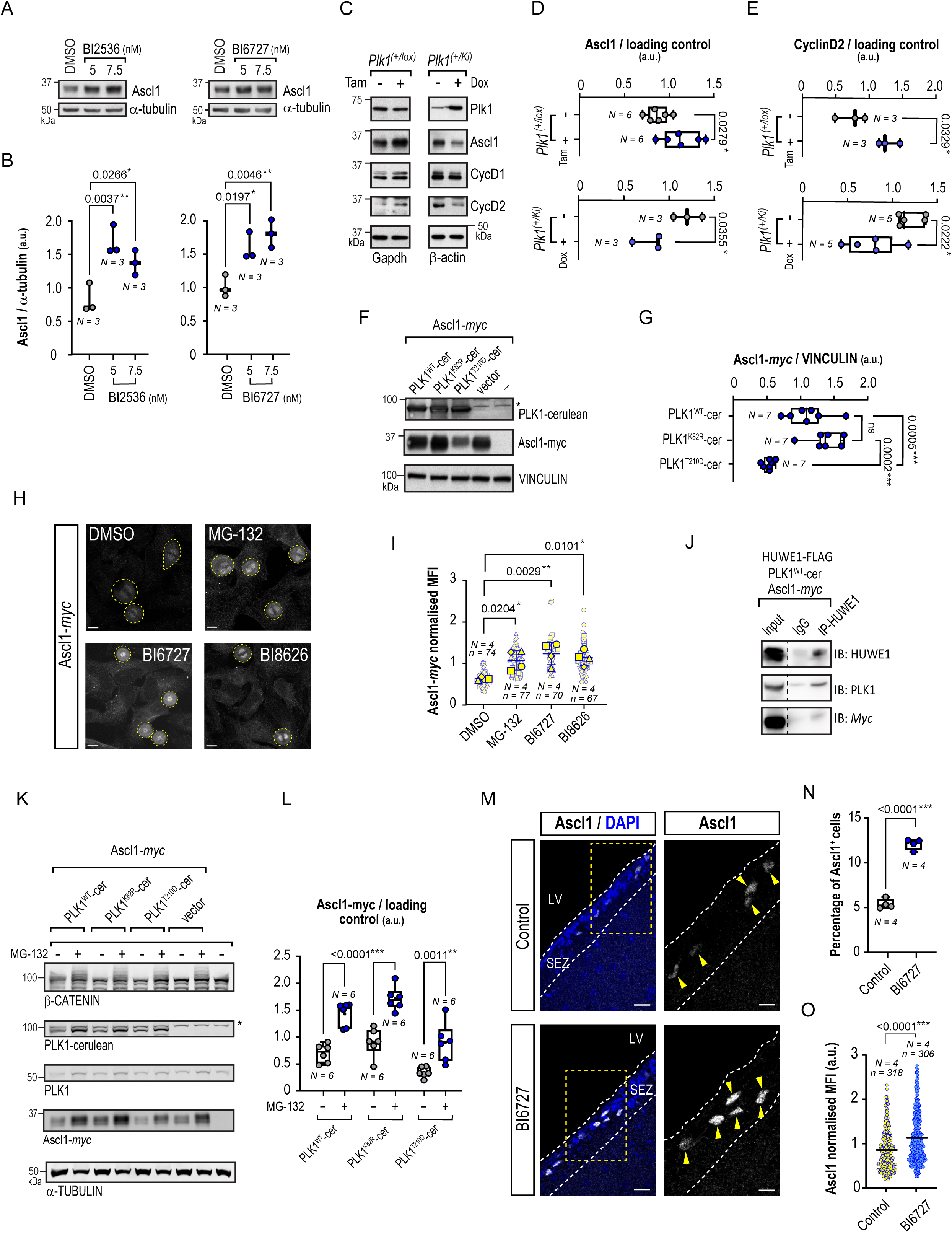
Plk1 functions upstream of the HUWE1-dependent regulation of Ascl1. (**A**) Representative immunoblot of Ascl1 and α-tubulin (loading control) in lysates from wild-type NSC-cultures treated with vehicle (DMSO) or the Plk1 inhibitors BI 2536 (left) and BI 6727 (right) at 5 and 7.5 nM. (**B**) Levels of Ascl1 normalized to loading control (α-tubulin) in control (DMSO) and BI 2536 (left) and BI 6727 (right)-treated wild type NSC protein extracts, represented in arbitrary units (a.u.) (N = 3, for each condition). (**C**) Representative immunoblots for Plk1, Ascl1, Cyclin D1, Cyclin D2 and Gapdh or β-actin for loading controls in *Plk1^(+/lox)^* and *Plk1^(+/K^*^i)^ NSCs in the absence or presence of 4-OH-tamoxifen (Tam) and Doxycycline (Dox), respectively. (**D)** Levels of Ascl1 in extracts from *Plk1^(+/lox)^* NSCs in the presence of vehicle (-) or Tam, and *Plk1^(+/K^*^i)^ NSCs in the presence of vehicle (-) or Dox, represented in arbitrary units (a.u.) after normalisation to the loading control (N = 6, *Plk1^(+/lox)^*; N = 3, *Plk1^(+/K^*^i)^ in both treatment conditions, respectively). (**E**) Levels of Cyclin D2 normalised to the loading control in extracts from *Plk1^(+/lox)^* NSCs in the presence of vehicle (-) or Tam (N = 3, in both treatment conditions), and *Plk1^(+/K^*^i)^ NSCs in the presence of vehicle (-) or Dox (N = 5, in both treatment conditions), represented in arbitrary units (a.u.). (**F**) Representative immunoblots of the protein levels of PLK1-cerulean, Ascl1-myc and VINCULIN (loading control) in lysates obtained from HEK293T cells co-transfected with *myc*-tagged *Ascl1* and either wild type human version of *PLK1-cerulean* (PLK1^WT^-cer), its kinase-dead mutant (PLK1^K82R^-cer) or a constitutively active mutant (PLK1^T210D^-cer). Cells co-transfected with an empty vector (vector) and untransfected cells (-) were used as controls. The asterisk indicates a non-specific band detected by the Plk1 antibody at 110 kDa. (**G**) Densitometric quantification of Ascl1-*myc* normalised to loading controls and expressed as arbitrary units (a.u.) in HEK293T cells co-transfected with Ascl1-*myc* and either PLK1^WT^-cer or mutants PLK1^K82R^-cer or PLK1^T210D^-cer. N = 7 independent transfections for each condition. (**H**) Representative confocal images of the immunodetection of Ascl1-*myc* (gray) in transfected HeLa cells in mitosis (yellow dashed lines) treated with MG-132, the Plk1 inhibitor BI 6727 or HUWE1 inhibitor BI 8626. Note the typical monopolar spindle-morphology provoked by the full inhibition of PLK1. (**I**) Mean fluorescence intensity (MFI) of Ascl1-*myc* in mitotic HeLa cells (n) treated with the different inhibitors (n = 74, 77, 70 and 67 independent cells, respectively) in N = 4 independent transfection experiments. (**J**) HUWE1 immunoprecipitation in HEK293T cells transfected with the indicated constructs, arrested in mitosis with nocodazole and treated with MG-132. A representative experiment showing interaction of HUWE1 with Plk1 and with Ascl1 is shown, out of three performed with similar results. Irrelevant lanes have been cropped out of the image (dashed black lines). (**K**) Representative immunoblots of PLK1-cerulean, PLK1 and Ascl1-*myc* protein levels obtained from untreated or MG-132-treated HEK293T cells co-transfected with *myc*-tagged *Ascl1* and either wild type *PLK1-cerulean* or its mutant versions. Cells co-transfected with an empty vector (vector) and untransfected (-) cells were used as controls. Alpha-TUBULIN levels are shown as loading control and β-CATENIN and higher molecular weight ubiquitylated forms are shown as control for proteasome inhibition. (**L**) Levels of Ascl1-*myc* normalised to the loading control represented in arbitrary units (a.u.), in extracts from untreated or MG-132-treated HEK293T cells, co-transfected with *Ascl1-myc* and either the wild type or mutant versions of *PLK1*. N = 6 for each condition. (**M**) Confocal micrographs of the staining for Ascl1 (gray) in control or BI 6727-treated wild-type mice. Cell nuclei were labelled with DAPI (blue). The white dashed lines delimit the SEZ. LV, lateral ventricle. Single channel images and insets (delimited by yellow dashed lines) for the staining of Ascl1 (right panels, gray, yellow arrowheads point at positive cells). Scale bars: 20 µm; inset: 10 µm. (**N**) Quantification of the percentage of Ascl1^+^ cells in the SEZ of control (N = 4) or BI 6727-treated (N = 4) wild-type mice. (**O**) Normalized mean fluorescence intensity (MFI) of Ascl1 *per* cell (n) in the SEZ of vehicle- (control; n = 318) or BI 6727-treated (n = 306) mice (N; N = 4 each). *P< 0.05, **p< 0.01 and ***p < 0.001, one-way ANOVA with *Dunnett’s* multiple comparison test (**B, G, I**), two-tailed unpaired Student’s *t*-tests (**D, E, N**), two-way ANOVA followed by *Šidák’s* multiple comparisons test (**L**) and Mann-Whitney’s test (**O**). Data are represented as box and whiskers from min. to max and all data points are shown (**B**, **D, E, G, L N** and **O**). Symbols represent mean values from independent neurosphere cultures obtained from individual mice (**B**, **D, E**), independent transfection experiments (**G, L**), independent mice (**N**) or individual cells (**O**). In (**I**) small symbols represent individual cells (*n*), and bigger symbols represent the mean values of each independent transfection experiment (*N*) and combined in a *superplot* representation. For full *Western* blot images refer to Fig. S7.

The degradation of Ascl1 by the proteasome mediated by Huwe1 occurs during mitosis (Liu et al., 2024), precisely when Plk1 expression and activity is at its highest (de Cárcer, 2019). To assess whether inhibiting Plk1 could provoke the stabilization of Ascl1 in mitosis we quantified the specific signal for *myc*-tag in mitotic HeLa cells transfected with *Ascl1-myc* (Fig. 3H). We found that BI 6727 induced an accumulation of *Ascl1-myc* comparable to that observed when the cells were treated with the proteasome inhibitor MG-132 or with Huwe1 inhibitor, BI 8626 (Fig. 3I). Moreover, we observed that in transfected cells with the human versions of both the kinase and the ligase and *myc*-tagged Ascl1, arrested in mitosis and treated with the proteasome inhibitor MG-132, Ascl1 and PLK1 co-immunoprecipitated with HUWE1 (Fig. 3J). Additionally, in PLK1 and Ascl1-*myc* immunoprecipitations we detected Ascl1-*myc* and HUWE1 or PLK1 and HUWE1 in both assays, respectively (Fig. S5), suggesting that the three proteins interact during this cell cycle phase. To further evaluate the functional association between Plk1 and Ascl1 *via* Huwe1, we tested whether its destabilization induced by Plk1 could be rescued by inhibiting the proteasome with MG-132. The treatment provoked the full rescue of Ascl1-*myc* levels (compared with untreated controls) in all the conditions tested, even in cells transfected with the constitutively active kinase (Fig. 3K-L). Altogether, our data indicate that Plk1 plays a role in targeting Ascl1 to degradation at the end of the cell cycle, upstream of the proteasome, likely via Huwe1.

Given the relevance of Huwe1/Ascl1 in controlling activation to quiescence transition in NSCs, we next set out to analyze Ascl1 levels in the SEZ of mice treated with BI 6727. Using immunohistochemical staining, we observed a two-fold increase in the percentage of Ascl1^+^ cells in the SEZ of BI 6727-treated mice vs. controls (Fig. 3M, N). Likewise, analyses of SEZ sections from *Plk1^Gfap^*^(+/Δ)^ *vs*. control mice revealed a significant increase in the percentage of positive cells immunostained for Ascl1 in Plk1-deficient mice (Fig. S5A-C). Next, we observed an increase in the levels of Ascl1 *per* cell in BI 6727 *vs*. control mice (Fig. 3O, Fig. S5D), which indicated an accumulation of Ascl1 protein also *in vivo*. Taken together, our results suggested that Plk1 functions to re-establish the adequate levels of the pro-activation factor Ascl1 at the end of the cell cycle thus serving to control the likeliness of NSCs to re-engage in proliferation in a successive round.

### Plk1 is a druggable target to boost adult neurogenesis

To investigate whether Plk1 inhibition-induced NSC activation could translate along the SEZ lineage and promote adult neurogenesis, we performed immunostainings for the neuroblast marker DCX in coronal sections of the SEZ of BI 6727- or vehicle-injected mice (Fig. 4A). We observed a significant increase in the percentage of DCX^+^ cells in BI 6727-treated mice compared to their corresponding controls (Fig. 4B). To explore whether neuronal output mirrored the enhanced production of neuroblasts observed, we used CldU-retaining as a marker for new neurons in the granule layer of the OB three weeks after injecting the mice with the analogue (Fig. 4C). We found that in BI 6727-treated mice, the number of CldU^+^ newly born neurons *per* area was significantly increased in comparison to the controls (Fig. 4D-E).

**Figure 4.**
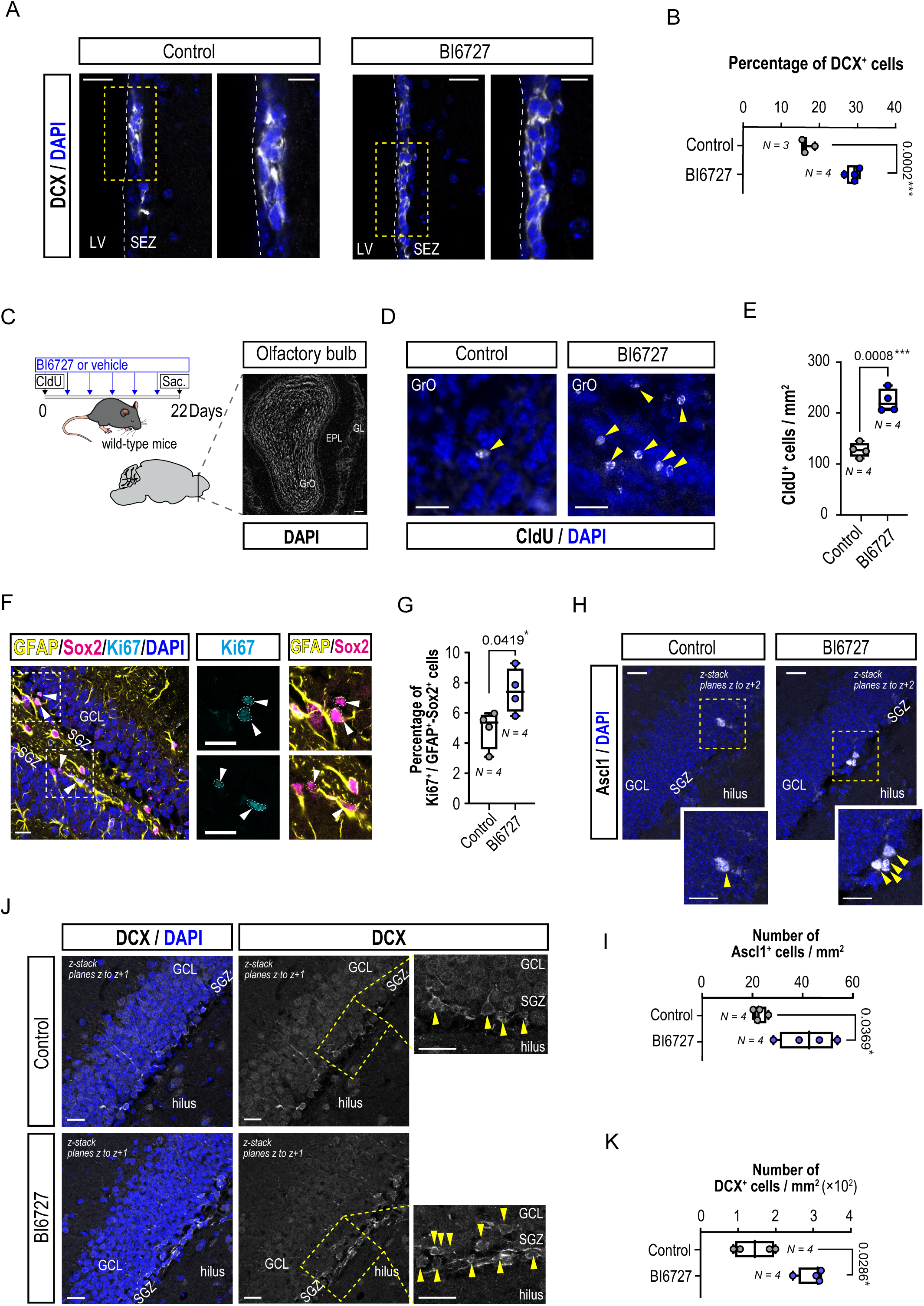
Plk1 is a druggable target to boost adult neurogenesis. (**A**) Representative images of coronal sections of the SEZ of control or BI 6727-treated mice immunostained for neuroblast marker doublecortin (DCX, gray). The white dashed lines delimit the boundary between the lateral ventricle (LV) and the SEZ. Cell nuclei were labelled with DAPI (blue). Scale bars: 20 μm; insets, yellow dashed lines: 10 μm. (**B**) Quantification of the percentage of DCX^+^ cells in the SEZ of control (N = 3) or BI 6727-treated mice (N = 4). **(C)** Schematic depicting the CldU and BI 6727 injection-chase protocol used in wild-type mice to label new-born cells in the olfactory bulb (OB), and representative image of an OB coronal section stained with DAPI (white) to visualize the different cell layers. GL, glomerular cell layer; EPL, external plexiform layer, GrO, granule cell layer. Scale bar: 100 μm (**D**) Representative images of the GrO of the OB of control or BI 6727 -treated mice immunostained for CldU (gray, yellow arrowheads). Nuclei are labelled with DAPI (blue). Scale bars: 25 μm. (**E**) Quantification of the total number of CldU^+^ cells *per* area in the GrO of the OB of control or BI 6727-treated mice (N = 4, each condition). (**F**) Representative images of coronal sections of the dentate gyrus (DG) of BI 6727-treated mice co-immunostained for GFAP (yellow), Sox2 (magenta) and Ki67 (cyan) in the subgranular zone (SGZ). GCL, granule cell layer. Cell nuclei were labelled with DAPI (blue). Scale bars: 20 μm; insets, white dashed lines: 20 μm. White arrowheads point at triple positive cells. (**G**) Quantification of the percentage of Ki67^+^ cells in the GFAP^+^-Sox2^+^ radial glia (RG)-like population in the DG of control (N = 4) or BI 6727-treated mice (N = 4). (**H**) Representative images of coronal sections of the DG of control and BI 6727-treated mice immunostained for Ascl1 (gray). Cell nuclei were labelled with DAPI (blue). Scale bars: 20 μm; insets, yellow dashed lines: 10 μm. Yellow arrowheads point at positive cells. (**I**) Quantification of the number of Ascl1^+^ cells in the DG of control (N = 4) or BI 6727-treated mice (N = 4). (**J**) Representative images of coronal sections of the DG of control and BI 6727-treated mice immunostained for DCX (gray). Cell nuclei were labelled with DAPI (blue). Scale bars: 20 μm; insets, yellow dashed lines: 10 μm. Yellow arrowheads point at positive cells. (**K**) Quantification of the number of DCX^+^ cells in the DG of control (N = 4) or BI 6727-treated mice (N = 4). Data are represented as box and whiskers from min. to max and all data points are shown (**B, E, G, I and K**). Dots represent mean values from independent mice (N), *P< 0.05 and ***p < 0.001, by two-tailed unpaired Student’s *t*-tests (**B, E, G and I**) and Mann-Whitney’s test (**K**).

We next sought to determine whether the other neurogenic niche, the SGZ of the DG was also susceptible to pharmacological stimulation by Plk1 inhibition, by using an alternative approach. By the simultaneous use of the markers GFAP and Sox2 in combination with Ki67, we found that RG-like DG-stem cells were also more prone to cycle in BI 6727-treated mice (Fig. 4F, G). Likewise, we found that Plk1 acute inhibition provoked an increase in Ascl1^+^ cells in the SGZ (Fig. 4H, I), and an enhancement of the neuroblast production, seen with anti-DCX staining (Fig. 4J, K), altogether indicating that Plk1 is a feasible target to induce NSCs-reactivation and neurogenesis *in vivo*.

## DISCUSSION

The discovery of the proteolytic regulation of Ascl1 by Huwe1 (Urbán *et al*., 2016) was a breakthrough in the field since it partly solved the puzzle of how NSCs could re-acquire quiescence. Although implicit was the notion that the re-establishment of quiescence should happen once the cell had actually divided, the direct link with the end of the cell cycle remained elusive. In this work we have identified the mitotic kinase Plk1 as an intrinsic regulator of NSC re-activation in adult neurogenic niches, upstream of the molecular axis Huwe1/Ascl1. Our results indicate that Plk1 functions as a mechanistical link of the degradation of Ascl1 to mitosis, serving to promote a low-activation state after NSC division.

The canonical function of Plk1 as a cell cycle driver and its pattern of expression in actively cycling cells of the SEZ might suggest that inhibiting Plk1, even partially, would impede NSC full activation, as is the case for cell cycle regulators RingoA, and CDK6, that both regulate threshold-acquisition for activation in neural and hematopoietic stem cells, respectively (Gonzalez et al., 2023; Laurenti et al., 2015). We, however, found the opposite phenotype, revealing an unanticipated role for Plk1 in NSCs. To disclose the molecular mechanism in a scenario in which Plk1 inhibition was promoting activation of NSCs, we decided to use asynchronous heterozygous cells, and so our results drew a much wider picture of the role of Plk1, highlighting not only its well-established canonical mitotic functions, but also unrecognized roles. GO-terms such as ‘chromosomal region’, ‘mitotic spindle’ and ‘nuclear pore’ found in our analysis comprise already described targets of Plk1 such as Numa1, Pcm1 and Npm1, nucleoporins (Nups), Ranbp2 and 3 and Tpr, supporting the role of Plk1 in nuclear pore complex disassembly and breakdown of the nuclear envelope in mitosis (Pintard and Archambault, 2018). However, the biggest cluster obtained in the CC-GO analysis was the one related to ‘synapse components’, comprising microtubule associated proteins (Mapt, Macf, Map2 amongst others), in line with cluster ‘regulation of neurogenesis’, obtained in the BP-GO analysis, in which these were also found. This was not unexpected, given the role of Plk1 in microtubule dynamics and spindle assembly, but also the cellular setting in which the analysis was performed. Intriguingly, novel functions of the kinase have been found in neurons in pathological contexts such as Amyotrophic Lateral Sclerosis, Alzheimer’s and Huntington’s diseases (Song et al., 2011; Szewczyk et al., 2025; Yamanishi et al., 2017). Thus, our data build on this idea and provide a starting point to investigate novel targets of Plk1 in the nervous system. Huwe1 participates in neuronal differentiation, DNA repair and apoptosis (reviewed in (Gong et al., 2020; Qi et al., 2022)) apart from its pivotal role controlling NSC quiescence degrading Ascl1. Herein, we provide evidence of the physical interaction between Plk1, Ascl1 and Huwe1, showing that the three proteins can be found in a complex during mitosis. The polo binding domain (PBD) of Plk1 binds to peptides phosphorylated either by a priming S/T-P kinase, or by Plk1 itself (self-priming), and interacts directly with Plk1’s substrates or alternatively, can act as tethers that recruit Plk1 to specific subcellular localizations (Zitouni et al., 2014). Ascl1 can be phosphorylated in SP sites by CDKs, ERK, and other proline-directed kinases (Ali et al., 2014; Ali et al., 2020; Azzarelli et al., 2022; Li et al., 2014; Liu *et al*., 2024; Wylie et al., 2015), and whereas it does not present a Plk1 consensus site, there is a putative PBD-site in Ascl1’s sequence (-S(pS/pT)P-, mAscl1^S88^/ hASCL1^S93^). In human cells PLK1 has been recently described as a general controller of the degradation of more than 200 substrates of E3 ubiquitin ligases SCF^βTrCP^ and SCF^CyclinF^ at G2/M (Mouery et al., 2024), but while substrates of βTrCP require phospho-degrons, Cyclin F substrates’ proteolysis seems to be independent of direct phosphorylation by PLK1. In the case of Atoh, another bHLH factor, a CK1-dependent phospho-degron is necessary for its ubiquitylation by Huwe1 (Cheng et al., 2016), and interestingly, the phosphorylation of Ascl1 by CDKs reduces its half-life (Azzarelli et al., 2024), which suggests the requirement of a phospho-degron, but however the full mechanism remains unexplored. Additionally to regulatory roles, Plk1 can play structural functions: it contributes to the epigenetic maintenance of centromeres by binding sequentially to Mis18 complex and HJURP, and this interaction is required for CENP-A deposition (Conti et al., 2024; Parashara et al., 2024). Hence, it is plausible that the binding of phosphorylated Ascl1 to Plk1 *via* PBD may function as a docking-site and facilitate its ubiquitylation by Huwe1. Whether the promotion of Ascl1 degradation by the proteasome relays or not on the phosphorylation of Huwe1 by Plk1 remains unexplored, although our data strongly indicate a dependency on the kinase’s catalytic activity. Huwe1’s sequence displays a consensus site for Plk1 phosphorylation -(D/E/N)X(pS/pT)-ϕ- (X represents any amino acid and ϕ a hydrophobic residue) (Burkard et al., 2009; Santamaria et al., 2011) at S^1368^ (Oppermann *et al*., 2012) and we detected 1 differentially (S^1395^) and 2 hypothetically (S^1907^ and S^2917^) regulated phosphosites in our study, being the latter a reduced consensus site for Plk1. The dissection of the precise mechanism, including the putative Plk1-dependent phosphorylation sites of Huwe1 in NSCs, while not the aim of this study, deserves further investigation.

*Ascl1* mRNA is regulated by the *Hes* family genes of the Notch effectors (Imayoshi et al., 2013; Sueda et al., 2019), and is induced in pNSC upon receiving activation signals, where it promotes its full activation (Andersen et al., 2014). Its degradation by Huwe1 ensures that a population of NSCs transits back into dormancy to avoid long-term exhaustion of the pool and depletion of the neurogenic potential of the niche (Blomfield et al., 2019; Urbán *et al*., 2016; Viñals et al., 2004; Xu et al., 2024). Although SEZ NSCs can repopulate by symmetric cell division (Obernier *et al*., 2018), the burst of NSC activation seen in young niches gradually declines, increasing quiescence to protect the population from self-consuming divisions (Encinas et al., 2011; Kippin et al., 2005; Pilz et al., 2018). Interestingly, a natural mechanism to prevent a NSC run out involves increasing the levels of Huwe1 to promote quiescence as ageing proceeds, thus safeguarding the neurogenic potential (Harris et al., 2021). SEZ NSCs can reactivate after self-renewal, a phenomenon that can occur right-away or with a lag of up to several months (Obernier *et al*., 2018). However, the link between the end of cell division and the re-establishment of quiescence *vs*. re-engagement in cell cycle was unknown, and this work has shed some light on this aspect. Our data indicate that Plk1 activity promotes the reacquisition of a more quiescent state in the daughter cells as part of the mechanism that degrades Ascl1 *via* Huwe1, during cell division. We speculate that the precise levels of active Plk1 will determine if the cellular offspring will inherit more Ascl1 protein, likely making one or both daughters more prone to reactivation in the next round. When *Ascl1* levels are artificially increased by *Hes* silencing, its sustained expression provokes a boost in neurogenesis (Sueda *et al*., 2019), as does Plk1 partial loss of function or graded inhibition. Additionally, our finding that Plk1 is a novel druggable target to boost NSC activation and neurogenesis both in the SEZ and the SGZ, has important implications for translatability. Evidence indicate the persistence of NSCs in the human brain however they become progressively inactive after infancy (Baig et al., 2024; Donega et al., 2019; Donega et al., 2022; Dumitru et al., 2025; Sanai et al., 2004). Nevertheless, their latent regenerative potential makes them a valid target for regenerative therapeutical strategies (Doludda *et al*., 2026). Our work extends the current understanding of the molecular mechanisms of quiescence in mammalian NSCs and broadens the possibility of pharmacologically reactivating endogenous dormant reservoirs for their potential use in future therapeutic approaches by means of small molecule inhibitors already in use in clinical trials.

## METHODS

### Mice

Mice with a floxed *Plk1*-allele crossed with *Pol2a-Cre*ERT2 strain and inducible-PLK1-FLAG-overexpressing mice were described previously (de Carcer *et al*., 2018; de Carcer et al., 2017; Trakala et al., 2015; Wachowicz *et al*., 2016). To conditionally inactivate Plk1 in NSCs, male *Plk1^(lox/lox)^* mice were crossed with females expressing Cre recombinase downstream of the murine glial fibrillary acidic protein (*Gfap*) promoter, obtained from Jackson Laboratories (Tg(Gfap-cre)73.12Mvs/J) (Garcia *et al*., 2004), to generate *Plk1^(+/lox)^;Gfap^(+/Cre)^* mice, and are referred herein to as *Plk1^Gfap(+/^Δ)*. *Plk1^(+/+)^;Gfap^(+/Cre)^* littermates were used as controls and are referred herein to as *Plk1^Gfap(+/+)^*. All mice were kept in a C57BL/6J (Jackson labs) background. Wild-type mice used were C57BL/6J. Two to 3 months-old and mature 8-month-old mice of both sexes were used. Mice were housed in a temperature-controlled environment, with a 12-hour light/dark cycle, and with water and food *ad libitum* at the Animal Facility Service of the Centro de Biología Molecular (CBM) Severo Ochoa under veterinary supervision and following the European Union 2010/63/EU guidelines and Spanish regulations RD-118/2021; ECC/566/2015. All procedures were approved by the Ethics Committee of Animal Experimentation of the CBM (UAM-CSIC) and local authorities. Genomic DNA was extracted from ear biopsies and used in PCR reactions for genotyping and confirmation of genomic recombination. Genotyping of fixed tissue sections was performed by PCR in DNA samples extracted using the standard protocol, followed by a 60 min incubation at 90 °C. Primers for all transgenes have been described elsewhere (de Carcer *et al*., 2018; de Carcer *et al*., 2017; Trakala *et al*., 2015; Wachowicz *et al*., 2016). For long-term label BrdU retention experiments, mice were injected and brains collected and processed, as described previously (Porlan et al., 2014). For the suboptimal pharmacological inhibition of Plk1 *in vivo*, 1.875 mg/kg Volasertib (BI 6727, Selleckchem) was administered by intraperitoneal injections in corn oil twice a week for three weeks.

### Antibodies

Information about the primary antibodies and dilutions used for the different applications is in Supplemental Spreadsheet 5.

### Immunohistochemistry

Immunostainings of the SEZ were performed in vibratome (Leica Microsistemas SLU, L’Hospitalet de Llobregat, Spain) free-floating sections, incubated for 48 h with primary antibodies (Supplemental Spreadsheet 5). The OBs were separated from the rest of the brain and sectioned with a cryostat (Leica Microsistemas). For the specific detection of BrdU or CldU, a previous 20 min 2N HCl denaturalization step was performed. For Ascl1 detection, the sections were subjected to heat-induced antigen retrieval in citrate buffer and a biotin/fluorescent-streptavidin incubations were used to amplify the signal. Immunofluorescent detections were performed with Alexa Fluor (Invitrogen) conjugated secondary antibodies. DAPI (1 μg/ml) was used for counterstaining. Samples were mounted using Fluoromount-G (SouthernBiotech). Images were acquired under the same settings for each experiment with LSM710 and LSM900 (Zeiss, Oberkochen, Germany) confocal laser scanning microscopes at the Optical and Advanced Microscopy Unit, CMB. Image acquisition was conducted serially throughout the area of the SEZ. At least 4 SEZs were photographed from each mouse. For OB neurogenesis the number of CldU^+^ cells present in the granular cell layer in at least 4 coronal sections of the medial region of the OB *per* mice was counted and normalized to the area measured in each sample. Quantitative analysis measurements were performed on the raw data with identical acquisition settings across samples. StarDist extension for ImageJ/Fiji software (v1.54f) was used for nuclei count to quantify Ascl1 pixel intensity in vivo. The binary masks of the nuclei of the SEZ cells were obtained, manually delimited following morphological criteria, and each cell was automatically identified as an independent region of interest (ROI) that were redirected to the original grayscale images. The MFI *per* cell of the marker Ascl1 was normalized to the mean value of all events. Postprocessing and analysis of the images was performed using ImageJ/Fiji software. For display of representative images, brightness/contrast of each channel was adjusted appropriately, and the same adjustments were applied consistently to the images being compared. Some images have been rotated and/or cropped for clarity.

### Cell culture

NSCs were obtained from the walls of the lateral ventricles of 2-3-month-old male and female mice and cultured as neurospheres (Belenguer et al., 2016) or in adhesive conditions using Geltrex (Invitrogen, Waltham, MA, USA,) as coating substrate, with murine natural EGF and human recombinant FGF-2 (both at 10 ng/ml) as mitogens, as previously described (Del Puerto *et al*., 2023). Primary neurosphere numbers were manually counted in an inverted microscope (Axiovert200, Zeiss ORCA-Flash4.0 LT sCMOS, Hamamatsu Photonics, Hamamatsu, Japan). Treatments used in NSC cultures were 1 µg/ml 4-OH-tamoxifen, 2.5 µg/ml doxycycline, and 5 and 7.5 nM BI 2536 (JS Research Chemicals Trading, Germany) and BI 6727 (Selleckchem, Houston, TX, USA), with the appropriate vehicle in each case (water, ethanol or DMSO). HEK 293T (ATCC, ref.: CRL-3216) and HeLa (ATCC, ref.: CCL-2) cells were cultured in DMEM supplemented with 10% FBS, 2 mM L-Glutamine and non-essential amino acids in a humidified atmosphere at 37°C and 5% CO_2_. All cultures were periodically tested for the absence of *Mycoplasma spp*. contamination. Transient transfection experiments were performed with Lipofectamine 2000 or Lipofectamine LTX Plus (both from Invitrogen) and cell lysates were obtained 48 h after transfection. Cells were treated with 40 μM nocodazole for 24 h, 2 μM MG-132 (both from Sigma-Aldrich) for 16 h prior to cell lysis or with 10 μM MG-132, 50 nM BI 2536 or 10 µM BI 8626 for 4 h prior fixation, depending on the assay. For immunostaining, HeLa cells were seeded on collagen treated coverslips, transfected and treated described and fixed using PFA. Cells were stained with anti-*myc* tag antibody and imaged in a LSM800 confocal microscope (Zeiss, Oberkochen, Germany), using the same laser settings within all samples. Cells undergoing mitosis were selected based on their chromatin condensation using DAPI. Maximum intensity projections (MIPs) were used to quantify the mean fluorescence intensity of Ascl1-*myc* in ImageJ/Fiji software.

### Plasmid constructs

pCer-C3-Plk1 (Addgene plasmid # 68132; http://n2t.net/addgene:68132; RRID:Addgene_68132), pCer-C3-Plk1-T210D (Addgene plasmid # 68133; http://n2t.net/addgene:68133; RRID:Addgene_68133), pCer-C3-Plk1-K82R pCer-C3-Plk1-K82R (Addgene plasmid # 68134; http://n2t.net/addgene:68134; RRID:Addgene_68134), all three, gifts from Catherine Lindon; pDARMO_CMVT_FLAG-HUWE1 (Hunkeler et al., 2021) was a gift from Eric Fischer (Addgene plasmid # 187121; http://n2t.net/addgene:187121; RRID:Addgene_187121). The sequence for *<u>myc</u>* <u>epitope</u> was inserted after the murine *Ascl1* sequence by PCR using the following primers (*Ascl1 F* 5’- GAA TTC GCC ACC ATG GAG AGC TCT GGC A -3’; *Ascl1-myc R* 5’- TCT AGA <u>CAG ATC TTC TTC AGA AAT AAG TTT TTG TTC</u> GAA CCA GTT GGT AAA GTC -3’). The resulting PCR product was cloned into the pGEM-T Easy vector (Promega) following the manufacturer’s instructions and subcloned in EcoRI / XbaI sites of pcDNA3 vector (Invitrogen).

### Label-Free Quantitative Phosphoproteomics Analysis

The analysis was conducted using a bottom-up strategy, originally incorporating four biological replicates for each condition. *Plk1^(+/lox)^*NSCs were seeded on Geltrex-coated plates and treated with 1 μM 4-OH-tamoxifen (or vehicle in the case of the controls) for 3 days prior to obtaining the protein lysates with SDS 1% buffer with protease- and phosphatase-inhibitors and lysates were snap frozen in dry ice. The procedure was performed at the proteomics facility of Centro Nacional de Biotecnología, as described previously in detail elsewhere (Martin-Hidalgo et al., 2020). The spectra were exported to *.mgf* format using Peak View v1.2.0.3 and analyzed using Mascot Server 2.6.1, OMSSA 2.1.9, X!TANDEM Alanine 2017.2.1.4, and Myrimatch 2.2.140 against a composite target/decoy database constructed from sequences in the *Mus musculus* reference proteome (Uniprot Knowledgebase, March 2019), along with common contaminants. The search engines configuration and included modifications were as described (Martin-Hidalgo *et al*., 2020). Score distribution models were employed to calculate peptide-spectrum match *p*-values, and spectra filtered by a False Discovery Rate (FDR) ≤ 0.01 (peptide level) were selected for quantitative analysis. Approximately 15% of the lowest quality total signals and full data from one biological replicate were excluded prior to further analysis due to inadequate quality. Differential regulation was assessed in the remaining three replicates using linear models, and statistical significance was determined using *q*-values (FDR). All analyses were performed using software from Proteobotics (Madrid, Spain). For the volcano plot representation only phosphopeptides present in >1 replicate were included. For over representation analysis, we generated a list of 833 genes from the protein names and their corresponding q-values maintaining said filters, and in the case of more than one phosphopeptides corresponding to the same protein, we chose the one with the lowest q-value. An additional filtering step of Log2 (fold change) >1.0 was applied. For ORA, *clusterProfiler* R package (v. 4.0) in R (v4.3.3) was used. Maps were created using the Cytoscape plug-in EnrichmentMap (Shannon et al., 2003). An FDR *q*-value of 0.01 was used as a node cutoff, as well as a gene-set similarity filter of 0.375 to correct for redundancy. KEGG pathway analysis was performed using *Enrichr* (Chen et al., 2013).

### Protein extracts and immunoblots

NSCs were lysed in in SDS-buffer (0 mM Tris-HCl pH 8.0, 1% SDS, 1 mM EDTA), and HEK293T in RIPA buffer (25 mM Tris-HCl, pH 7.6, 1% Triton X-100, 0.5% sodium deoxycholate, 0.1% SDS, 150mM NaCl), with protease and phosphatase inhibitors at 4°C. Lysates were incubated on a rotary shaker for 30 min and cleared by centrifugation at 200 × g for 15 min at 4°C. Protein concentration was determined with BCA Protein Assay kit (Pierce Biotechnology Inc., Rockford, IL, USA). 20-50 µg of total protein were subjected to SDS-PAGE and transferred to nitrocellulose membranes (BioTrace NT, Pall Corporation, Pensacola FL, USA). Membranes were blocked in PBS or TBS containing 3% BSA or 5% skimmed milk, and 0.025% Tween-20 and incubated with primary antibodies (Supplemental Spreadsheet 5), followed by appropriate peroxidase-conjugated secondary IgGs (Dako Agilent, Santa Clara, CA, USA) before performing chemiluminescence reactions (Western Lightning Plus; Perkin Elmer, Waltham, MA, USA) and imaging. In some cases, a stripping protocol was used. Efficient removal of the previous signal was assessed prior to re-blotting the membranes. Both autoradiographic films and digital imagers (Amersham 600 or 680) were used across the study. Unsaturated signals were quantified using ImageJ/Fiji software, normalized to their loading controls and to the mean of all data *per* experiment and represented as arbitrary units. The respective band intensities were quantified with ImageJ/Fiji software using the raw images, either from digital imagers or from films scanned using a calibrated densitometer (BioRad). Values were normalised to the pertinent loading control and to the mean intensity value for each independent experiment and represented as arbitrary units. Immunoblot images have been cropped for presentation, and in some cases irrelevant lanes have been cropped out (indicated by a dashed line). All uncropped Western blot images are presented in Fig. S7.

### Coimmunoprecipitation

HEK293T cells transfected with Ascl1-*myc*, pCer-C3-Plk1 and pDARMO_CMVT_FLAG-HUWE1 were arrested in mitosis using 40 nM nocodazole for 24 h. Cells were lysed in 20 mM Tris-HCl pH 7.4, 100 mM NaCl, 0.5 mM EDTA, 0.1% NP-40 with protease and phosphatase inhibitors. 50 µg of clarified lysate was used as input, and 1 mg of total protein was incubated with primary antibodies (Supplemental Spreadsheet 5) or IgG isotype control for 16 h at 4 °C while rocking, prior incubation with 25 µl Dynabeads Protein G (Invitrogen) for 1 h at 4 °C. Samples were washed twice with 20 mM Tris-HCl pH 7.4, 100 mM NaCl, 0.5 mM EDTA and once with lysis buffer. Samples were boiled in Laemmli-sample buffer for 5 min and resolved by SDS-PAGE followed by Western blot as described above.

### Statistical Analyses

Analyses of significant differences between means were performed with Prism 9 software (GraphPad, San Diego, CA). Statistical tests used are specified in figure legends. Data were tested for normality and homogeneity of variances. Angular transformations were performed to meet test assumptions when needed. Welch’s correction or a non-parametric test were used when homoscedasticity or normality were not met, respectively. The number of biological replicates (N) carried out with independent subjects (mice, NSC-cultures established from different mice, independent transfections or cells) is shown in each bar graph as dots and is specified in figure legends. Outliers were identified and removed when applicable using the ROUT method. In all cases, p < 0.05 denoted statistical significance. ANOVA tables for Fig. 3B, 3G, 3I and 3L are provided in Supplemental Spreadsheet 6). Investigators were not blinded to allocation during the experiments, use of animals or outcome assessment in every case. For animal studies a stratified randomization was used to ensure similar age, and both sexes were used. Sample sizes were based on previously published experiments.

## ACKNOWLEDGMENTS

This work was funded by grants PID2023-146945NB-I00 to EP, PID2023-153284OB-100 to TI, PID2023-152962NB-I00 to IF, PID2024-161681OB-10 to MM, and RED2024-153635-T (iDIFFER) to IF and MM, funded by MCIU/AEI/ 10.13039/501100011033 and by ERDF/EU; TI, IF and MM are also funded by Centro de Investigación Biomédica en Red (CIBER) de Enfermedades Neurodegenerativas (CIBERNED, to TI and IF) and Cáncer (CIBERONC, to MM), Instituto de Salud Carlos III, Spain. IF is recipient of grants CIPROM/2024/57 from Generalitat Valenciana, and ERC Advanced Grant 101098241 from the European Commission. We are grateful to S. Benito, B. Ortigosa and B. Alcover for excellent technical support, to the Advanced Light Microscopy Facility (SMOA), and to the animal facility at the Centro de Biologia Molecular Severo Ochoa (CBM).

## AUTHOR CONTRIBUTIONS

ALB-M, CL-F, EG-G, JMM-R, BB-S, MA-C, FZ, and EP performed research. MM generated Plk1-floxed and iPLK1 mice. TI, IF and MM provided reagents and tools. All authors contributed to data analysis and interpretation and to the final version of the manuscript. EP performed study concept and design, secured funding, supervised the work and conceived and wrote the manuscript.

## SUPPLEMENTARY INFORMATION

**Figure S1.**
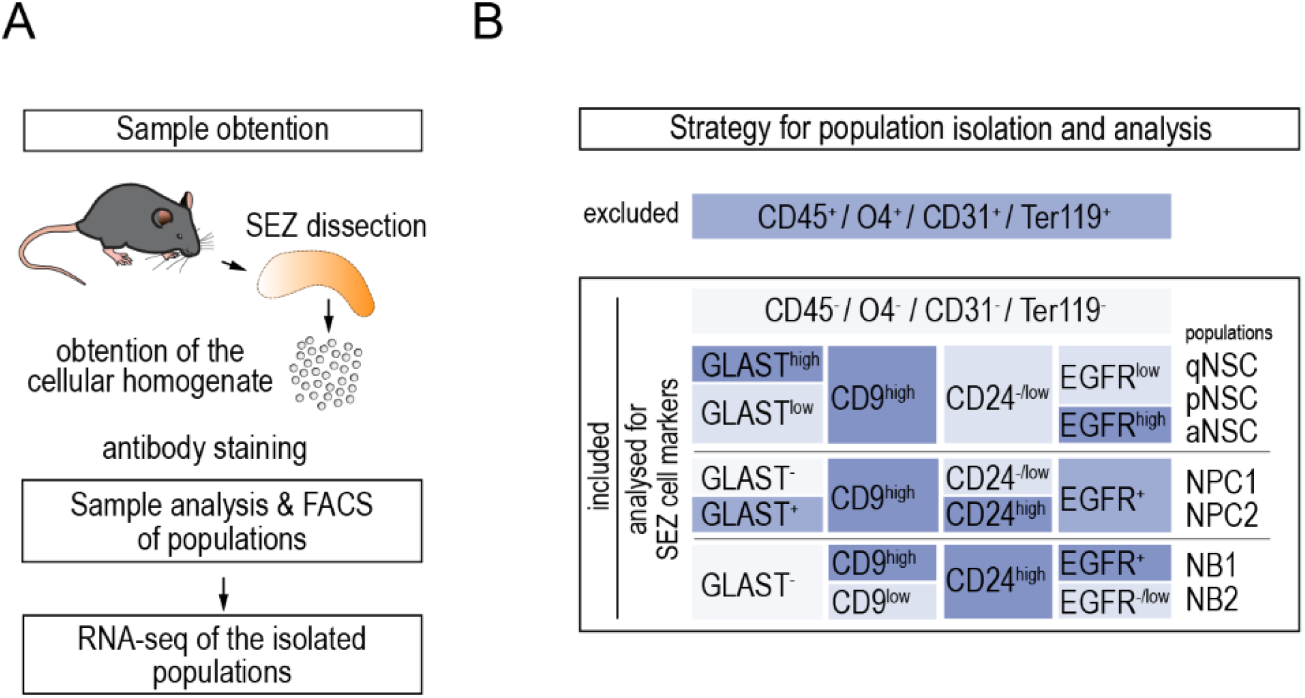
(**A**) Outline of the workflow followed in Belenguer et al., 2021 for the obtention of the samples. (**B**) Strategy and antibody panel used for the immunolabeling of SEZ cells and the prospective isolation of the different populations, quiescent (q), primed (p) and active (a) NSC, Neural Progenitor Cells 1 and 2 (NPC1, 2) and neuroblast population 1 and 2 (NB1, 2). Modified from Belenguer et al., 2021.

**Figure S2.**
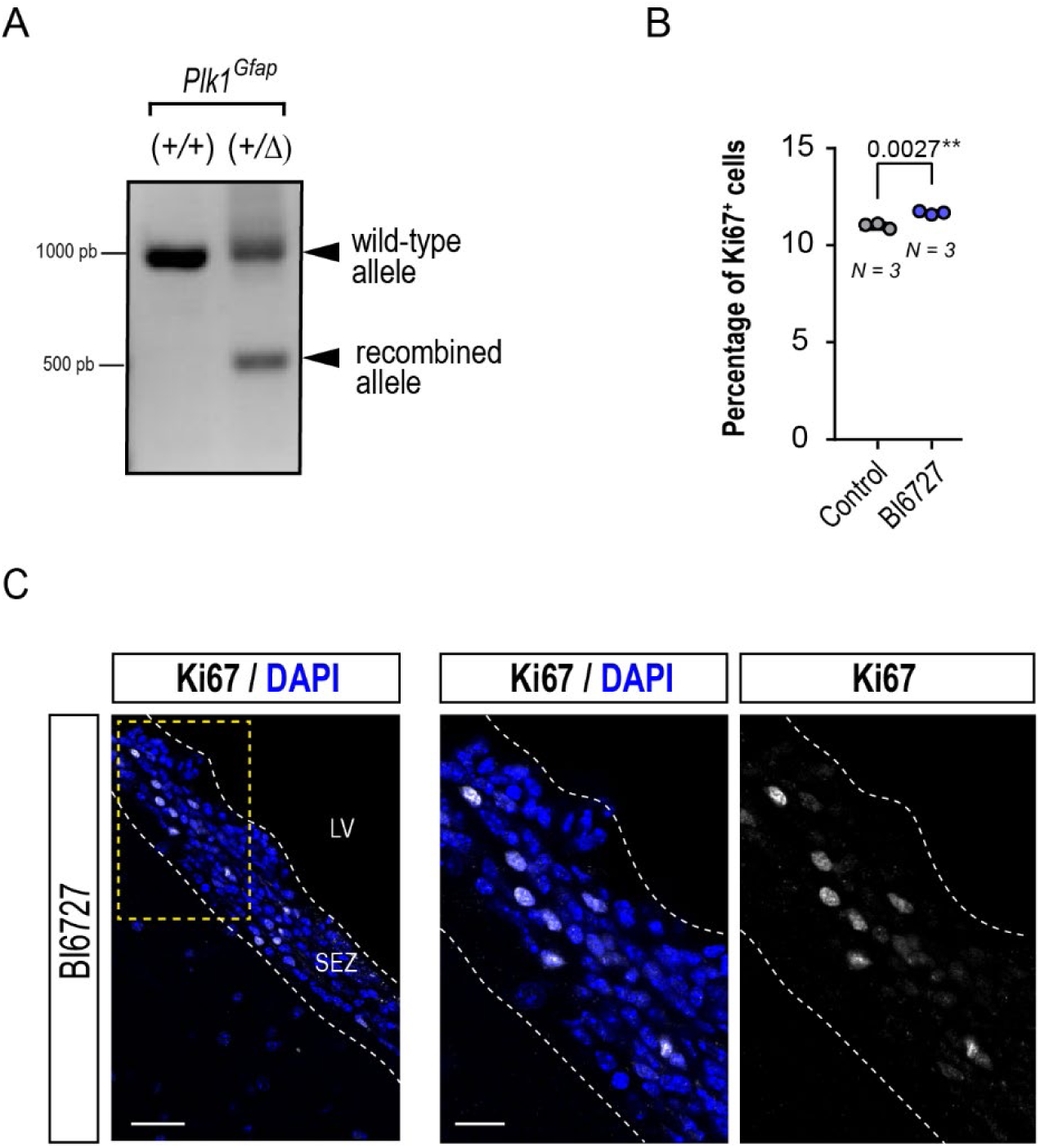
**(A)** Representative PCR for the *Plk1*-floxed recombined allele in brain tissue sections from *Plk1^Gfap(+/+)^* and *Plk1^Gfap(+/Δ)^*mice. (**B**), Percentage of Ki67^+^ cells in the SEZ of vehicle (control, N = 3) and BI 6727-treated (N = 3) mice. **(C)**, Representative confocal micrographs for Ki67 (gray) in BI 6727-treated mice. Scale bars: 50 μm, and 20 μm (inset). The white dashed lines delimit the boundary between the lateral ventricle (LV) and the subependymal zone (SEZ). Cell nuclei were labelled with DAPI (blue). Data are represented as box and whiskers from min. to max and all data points are shown. **p < 0.01, unpaired two tailed Student’s *t* test. Each symbol represents the mean value from an individual mouse (N).

**Figure S3.**
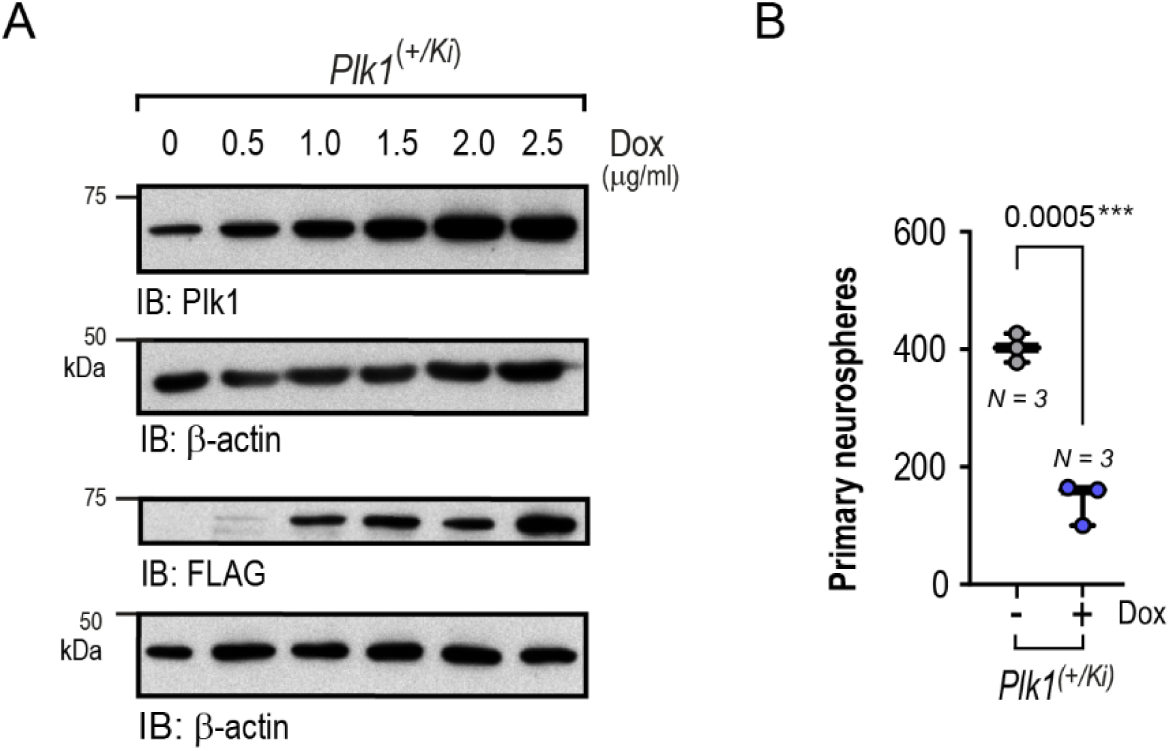
**(A)** Representative *Western* blots in lysates obtained from cultures of *Plk1^(+/Ki)^* NSCs treated with increasing doses of doxycycline (Dox) to induce the expression of a FLAG-tagged human *PLK1* transgene. Samples were run in parallel and detected with anti-FLAG or Plk1, and β-actin antibodies for loading control. (**B**) Quantification of the number of primary neurospheres obtained from *Plk1^(+/Ki)^* NSCs in the absence or presence of Dox. Values were normalized to vehicle-treated cells. N = 3 for both conditions. Data are represented as box and whiskers from min. to max and all data points are shown. \*\*\**p* < 0.001, two-tailed paired Student’s *t*-tests. Each symbol represents one independent neurosphere culture obtained from different mice (N).

**Figure S4.**
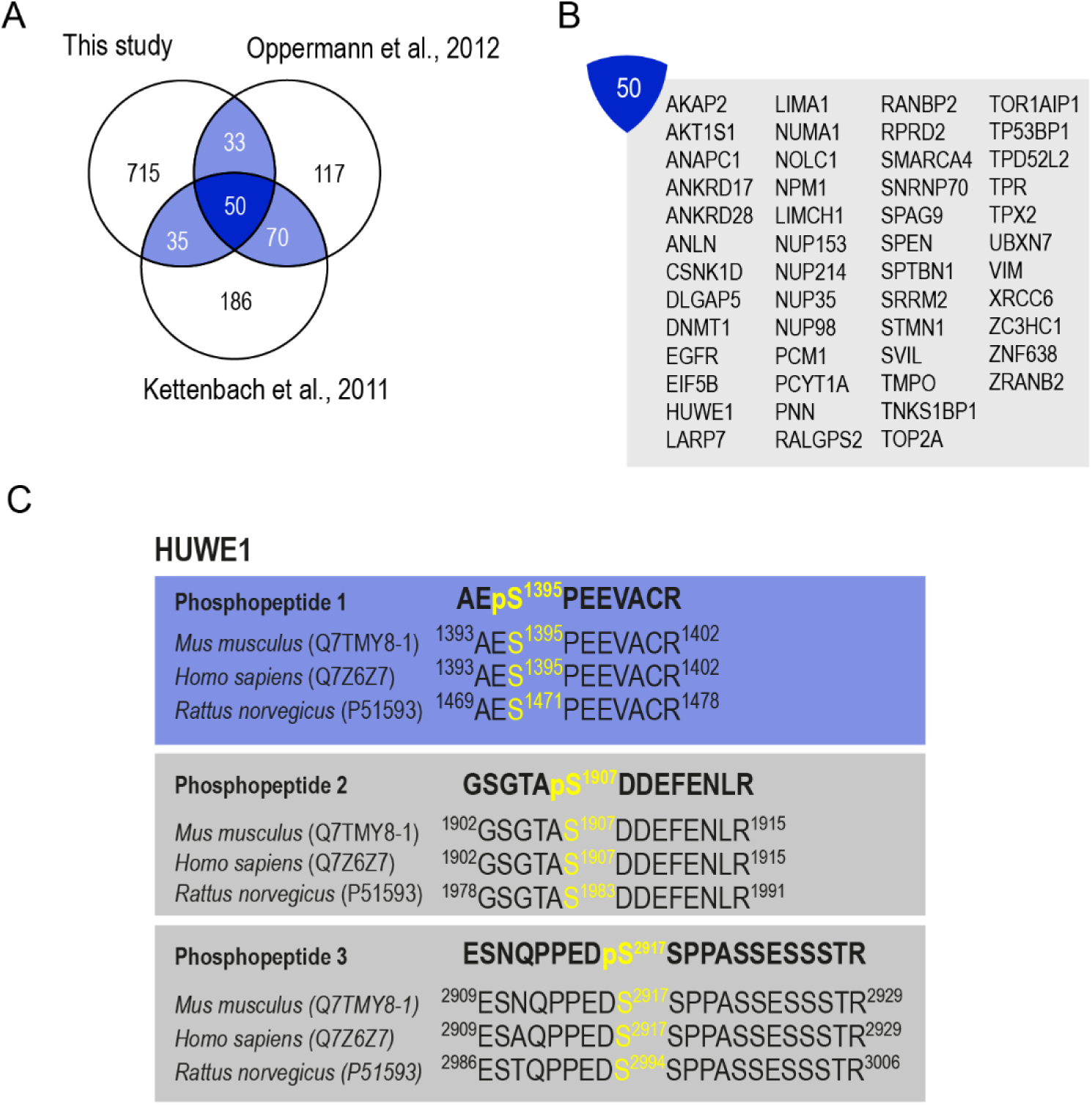
**(A)** Venn diagram indicating the number of proteins in this study common with studies of Oppermann et al., 2012 and Kettenbach et al., 2011. (**B**) List of 50 common proteins to the three studies. (**C**) HUWE1 phosphopeptides found in our study showing the modified residue (yellow). Blue box denotes that the phosphopeptide was differentially regulated (q < 0.05), whereas gray boxes denote a putative regulation (q > 0.05) in *Plk1^(+/Δ)^* NSCs.

**Figure S5.**
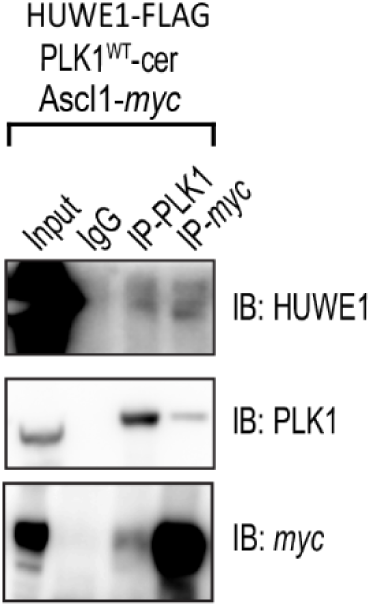
Immunoprecipitation (IP) of PLK1 and Myc, showing the reciprocal interaction between Ascl1 and PLK1, and their interaction with HUWE1 in mitosis HEW293T cells. Representative immunoblots (IB) for the specific detection of HUWE1, PLK1 and Ascl1-*myc* in samples immunoprecipitated for PLK1 and *Myc* in lysates from transfected cells. IgG, isotype control antibody. One representative experiment out of three performed with similar results is shown.

**Figure S6.**
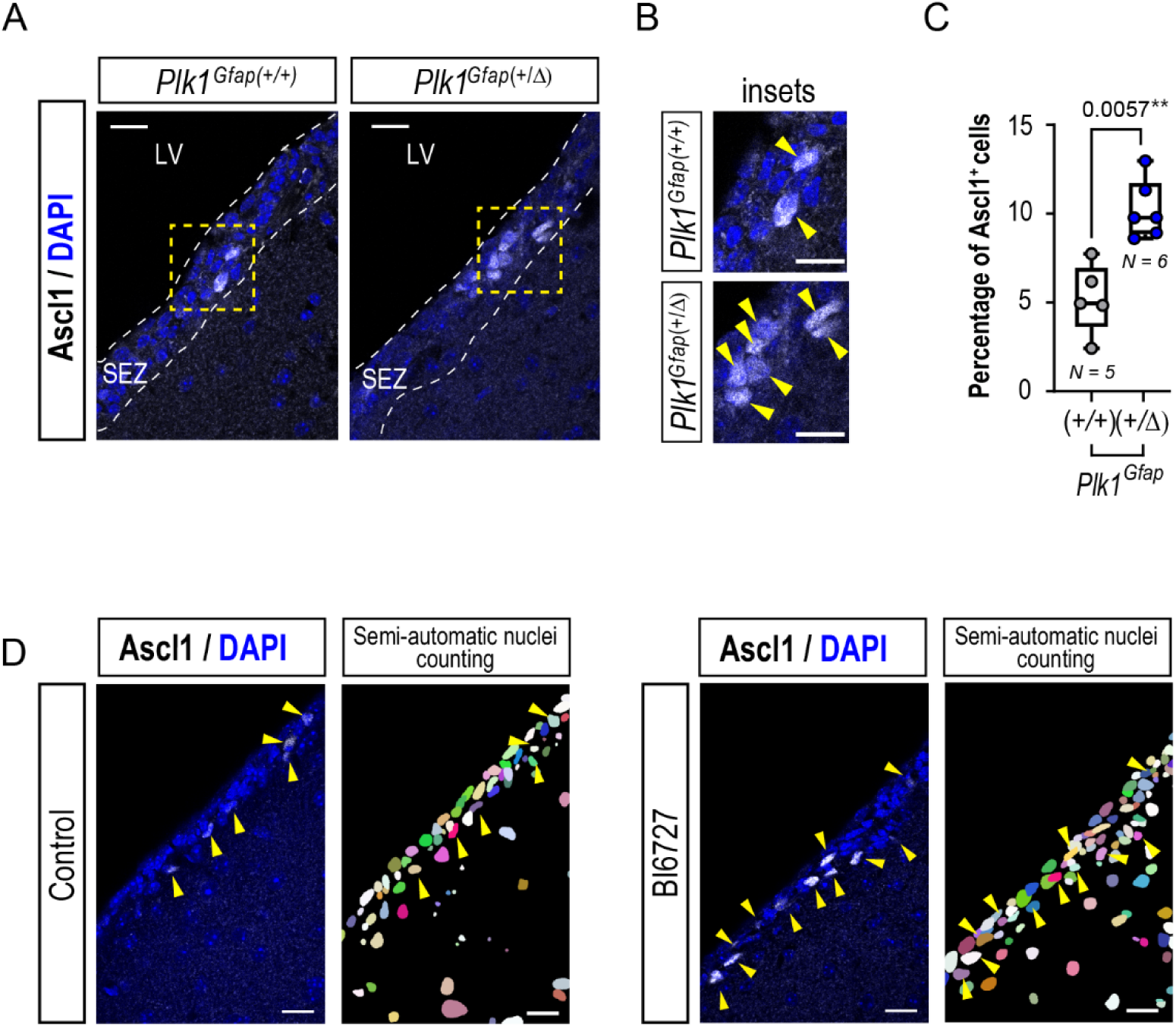
(**A**) Confocal micrographs of the staining for Ascl1 (gray, yellow arrowheads) in the SEZ (delimited by white dashed lines) of *Plk1^Gfap(+/+)^* and *Plk1^Gfap(+/^Δ)* mice. Nuclei are stained with DAPI (blue). LV, lateral ventricle. (**B**) Insets showing magnification of the delimited areas in (**A**) by yellow dashed squares. Scales: 20 μm and 10 μm (insets). **(C)** Quantification of the percentage of Ascl1^+^ cells in the SEZ of *Plk1^Gfap(+/+)^* (N = 5) and *Plk1^Gfap(+/^Δ)* (N = 6) relative to the total number of cells. Data are represented as box and whiskers from min. to max and all data points are shown. Each symbol (N) is the mean value from an individual mouse; **p < 0.01, unpaired two tailed Student’s *t* test. (D) Example of the segmentation mask used in bioimage analyses for the quantification of the signal of Ascl1 *in vivo* shown in Fig. 3A. Confocal micrographs of the staining for Ascl1 (left panels, gray, yellow arrowheads point at positive cells) in control or BI 6727-treated wild-type mice. Cell nuclei were labelled with DAPI (blue). Scale bars: 20 µm.

**Figure S7.**
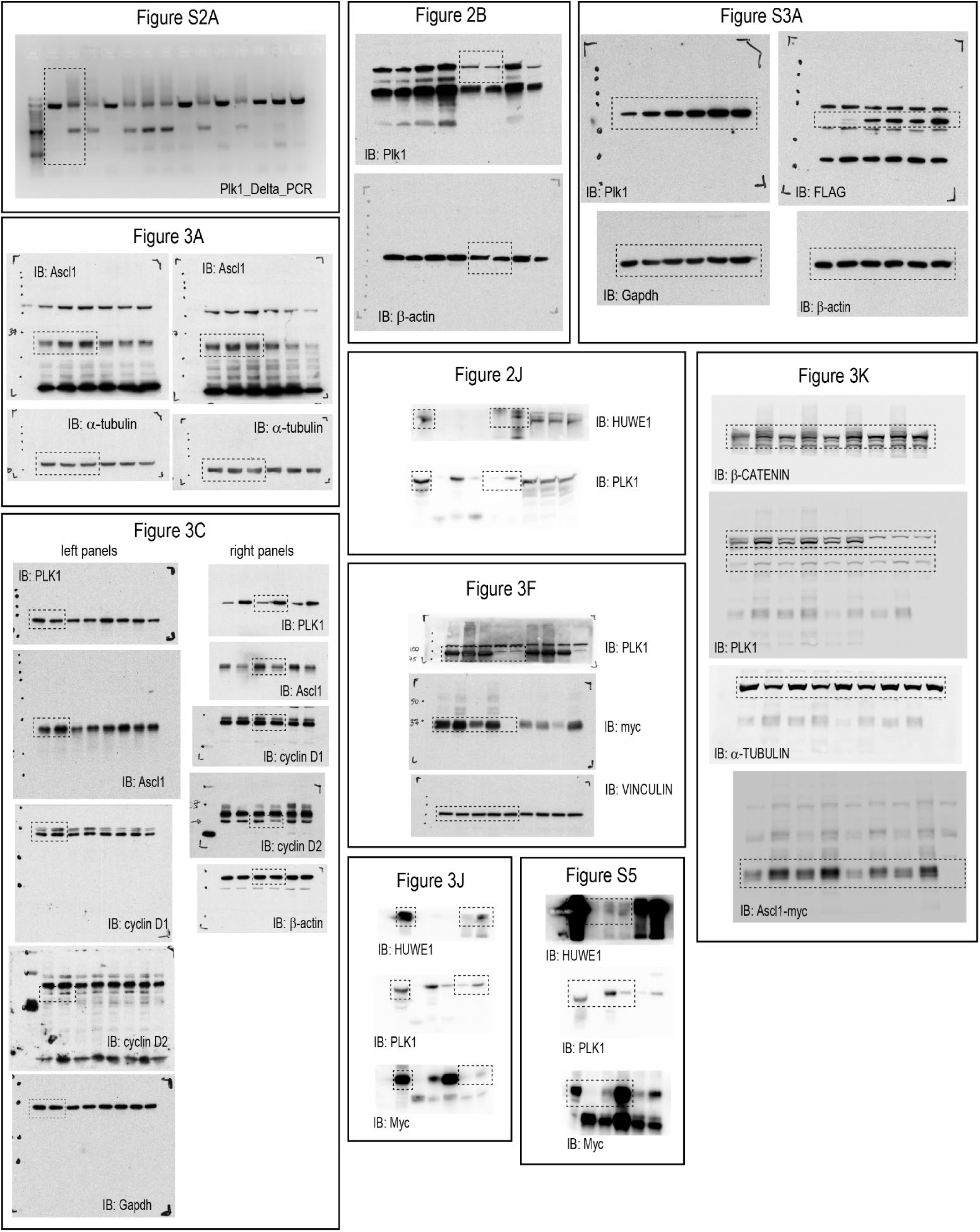
Uncropped gels and *Western* blots. IB, immunoblot.

Supplemental_Spreadsheet_1

data in this spreadsheet is deposited in https://doi.org/10.5281/zenodo.20036962

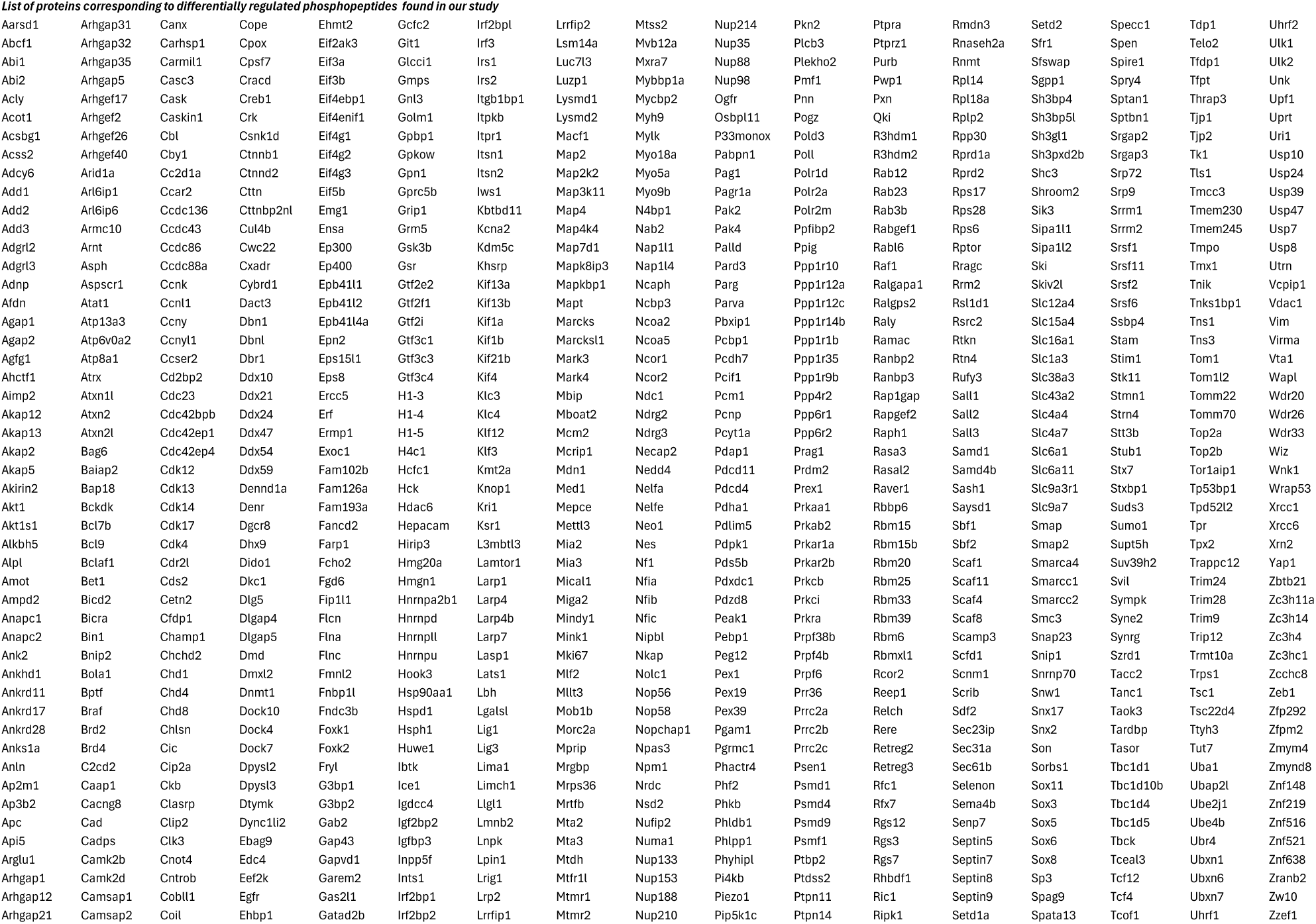
Supplemental_Spreadsheet_2.

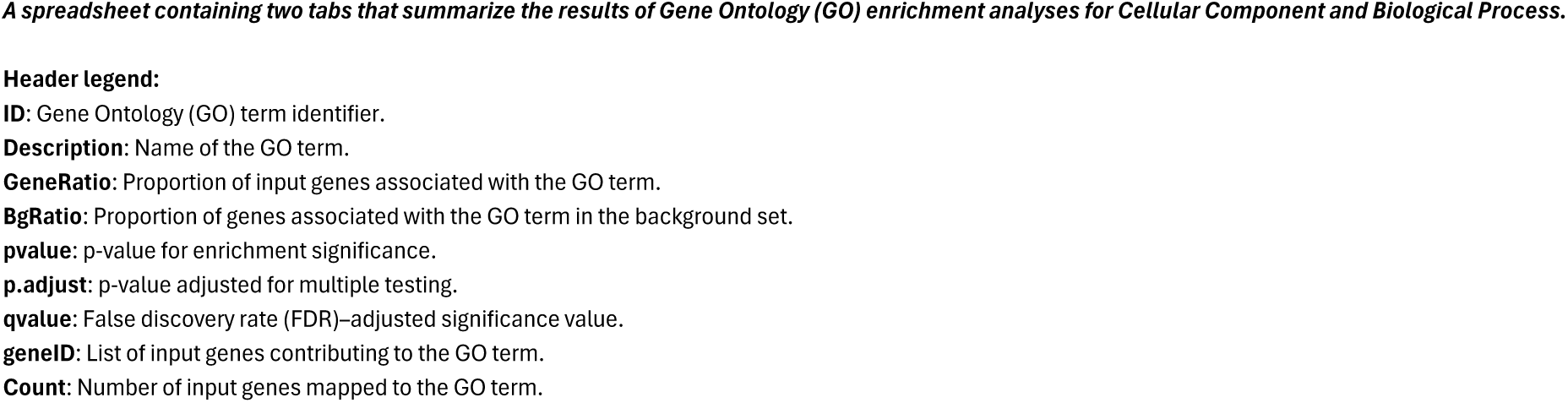
Spreadsheet_3_Tab_1_Legend.

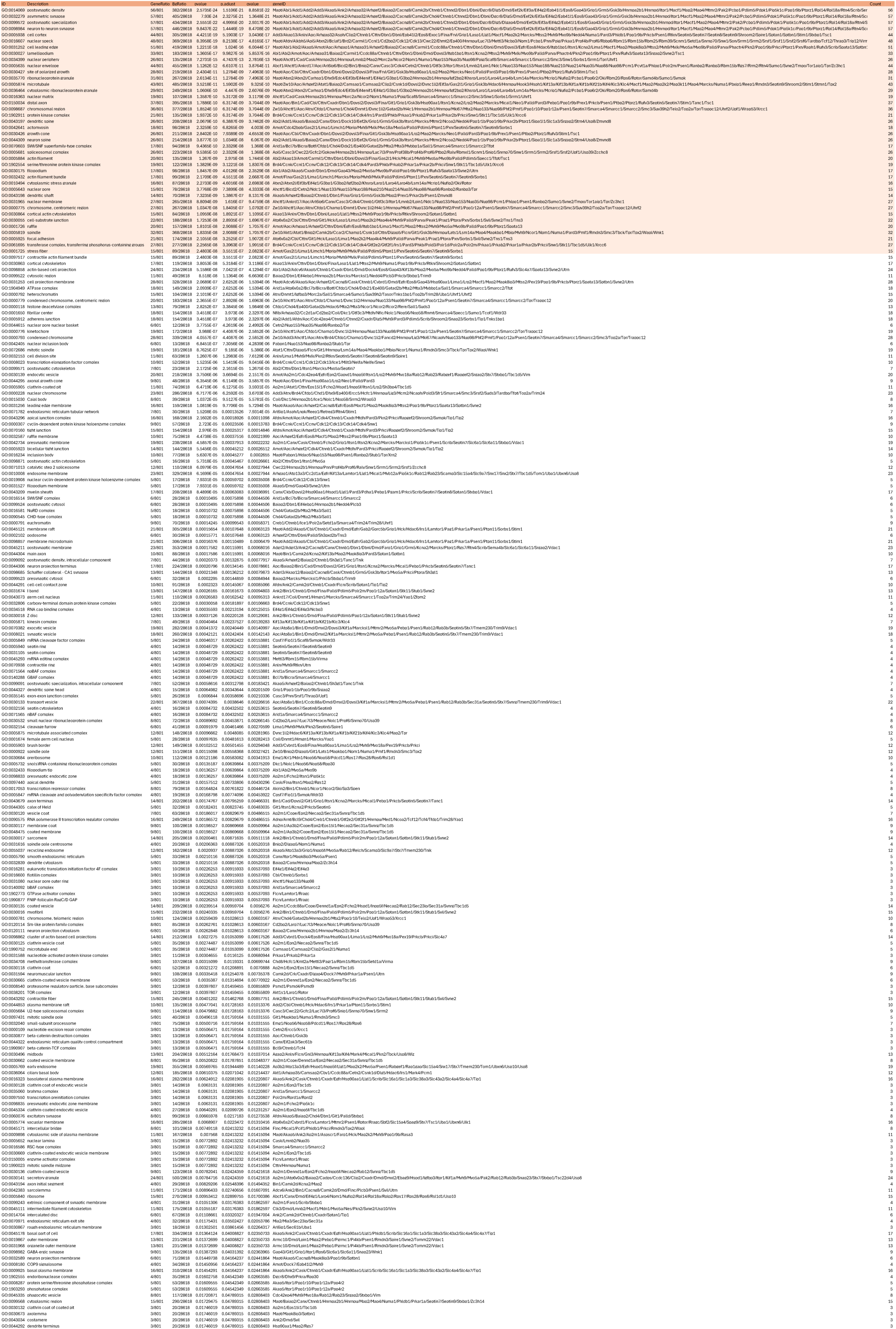
Spreadsheet_3_Tab_2.

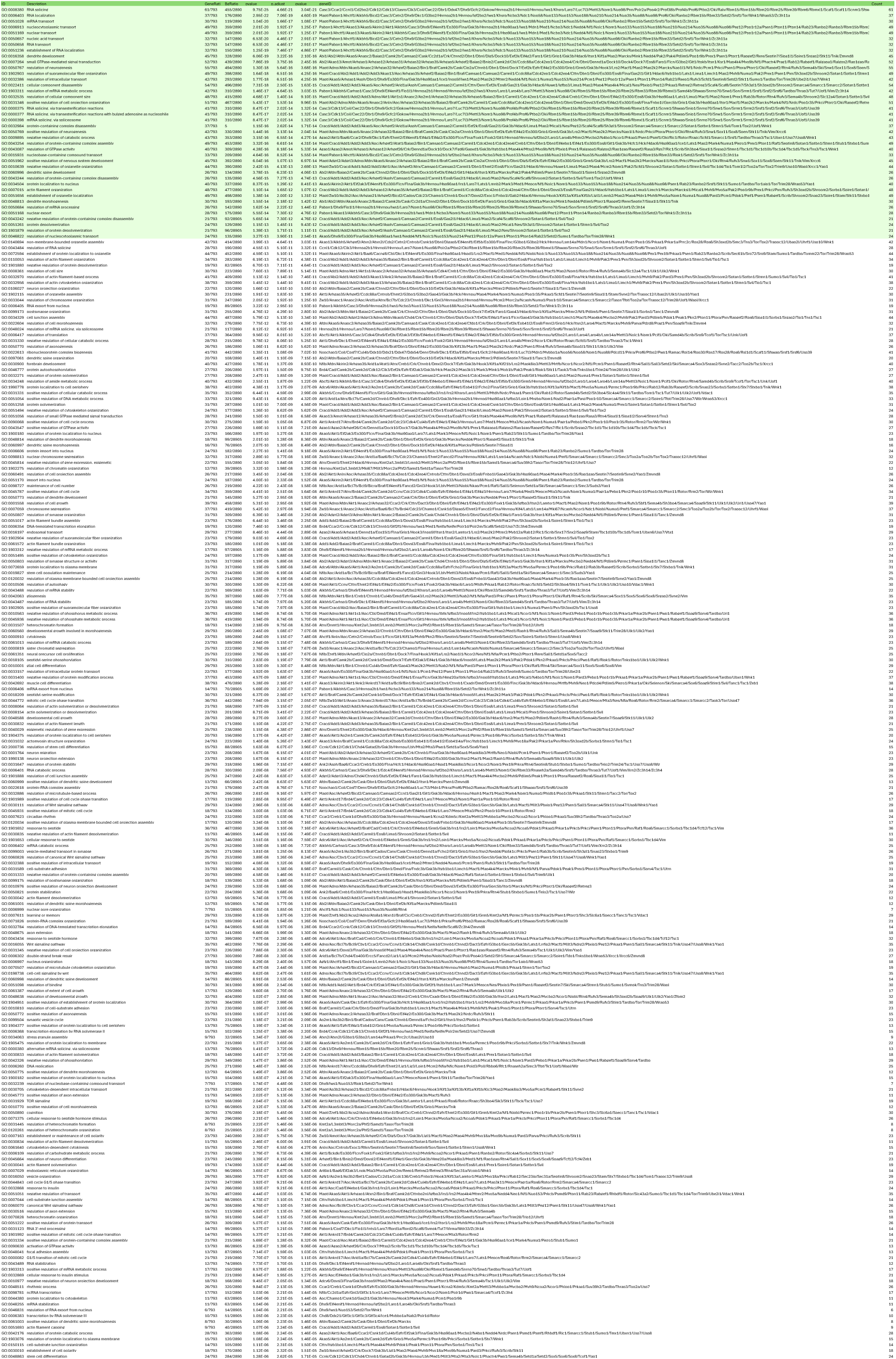
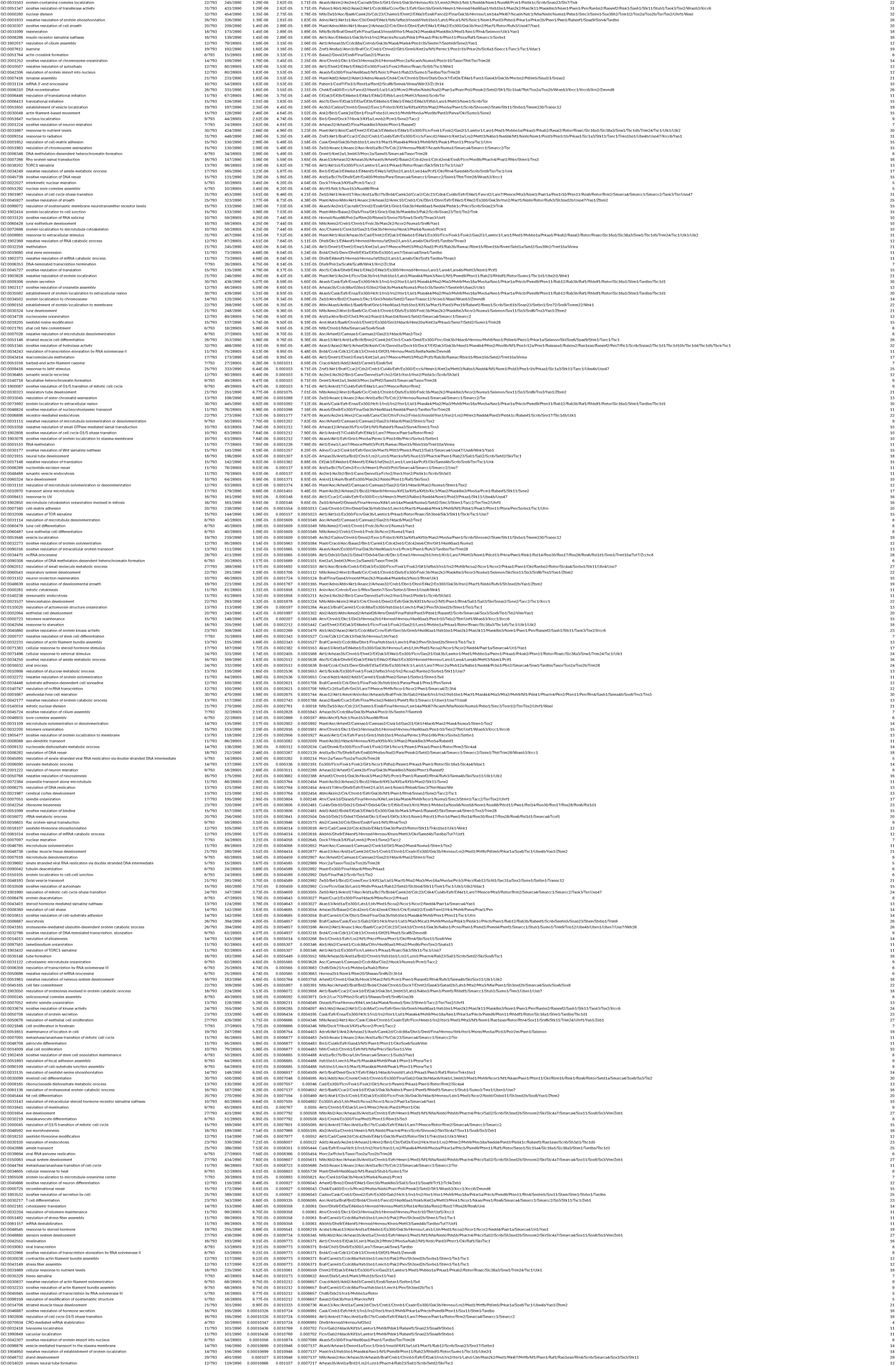
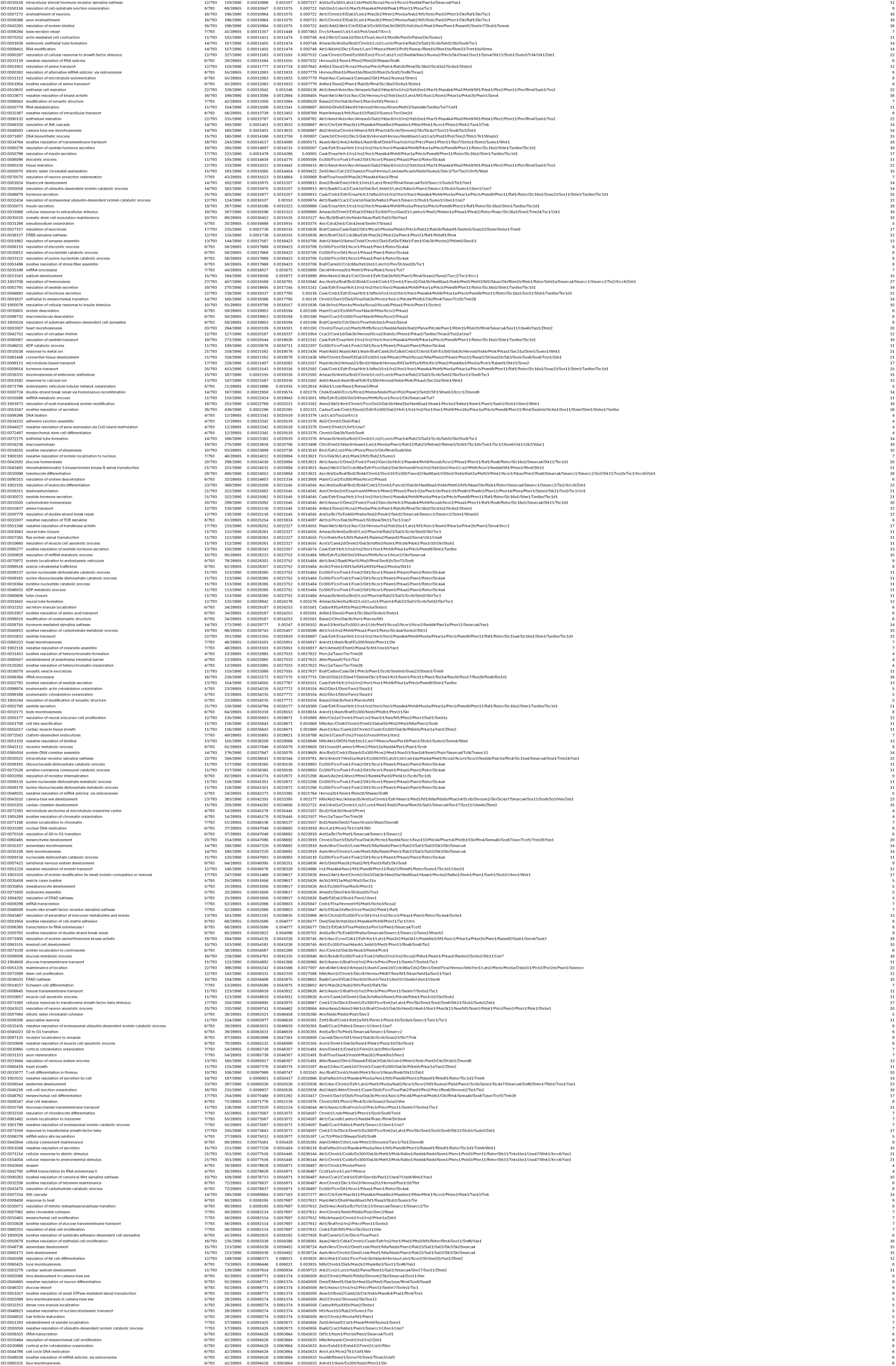
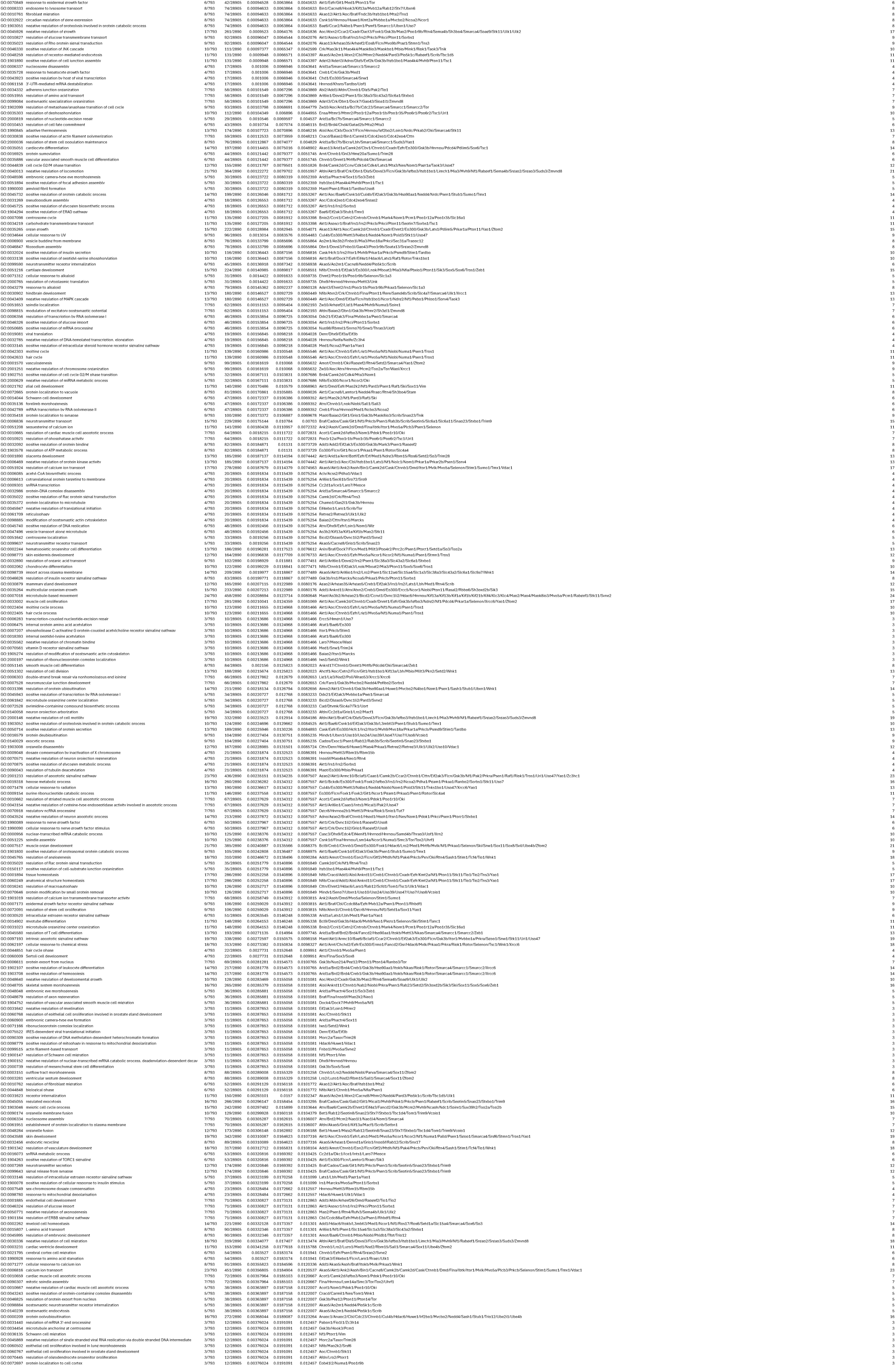
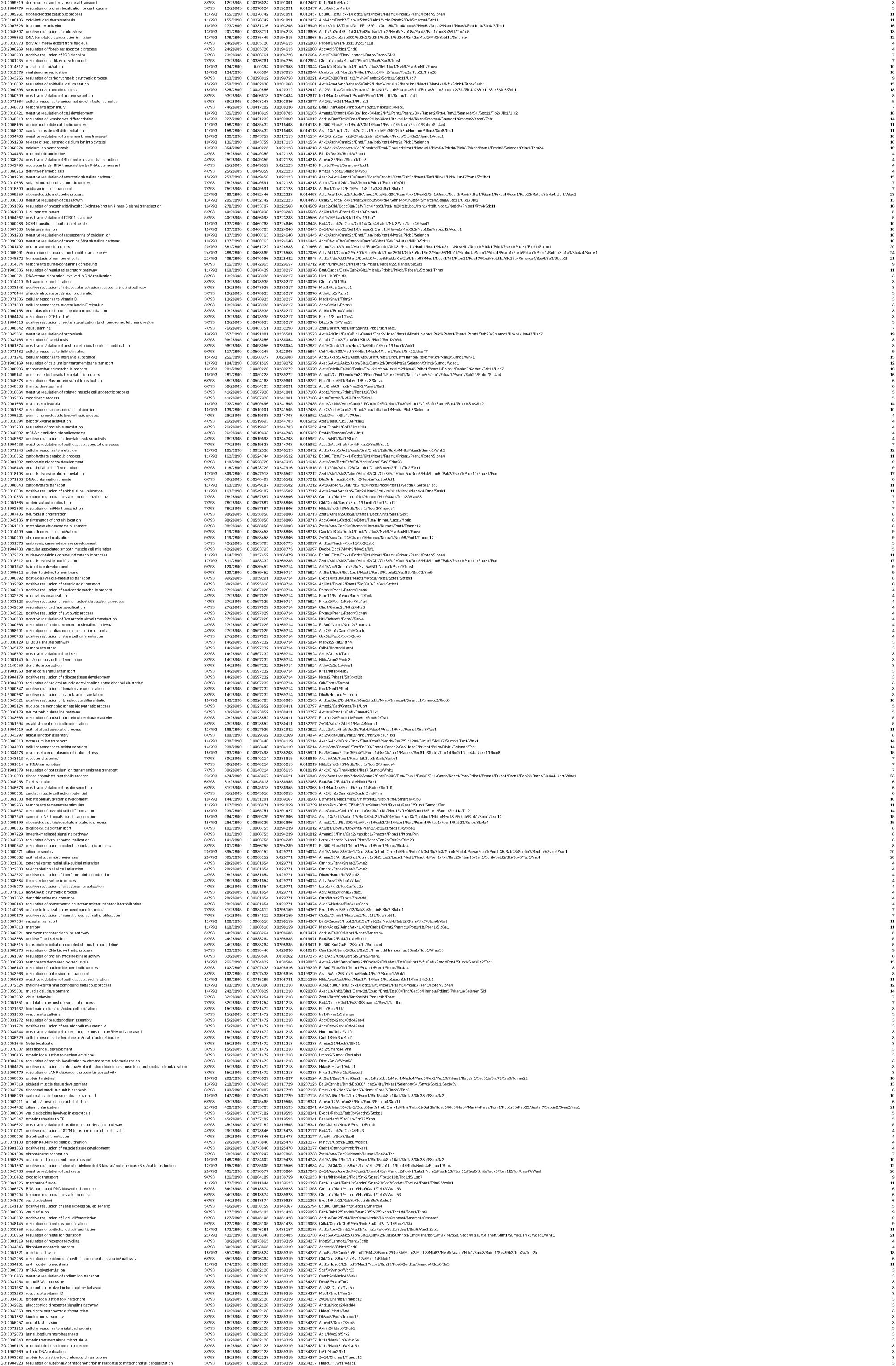
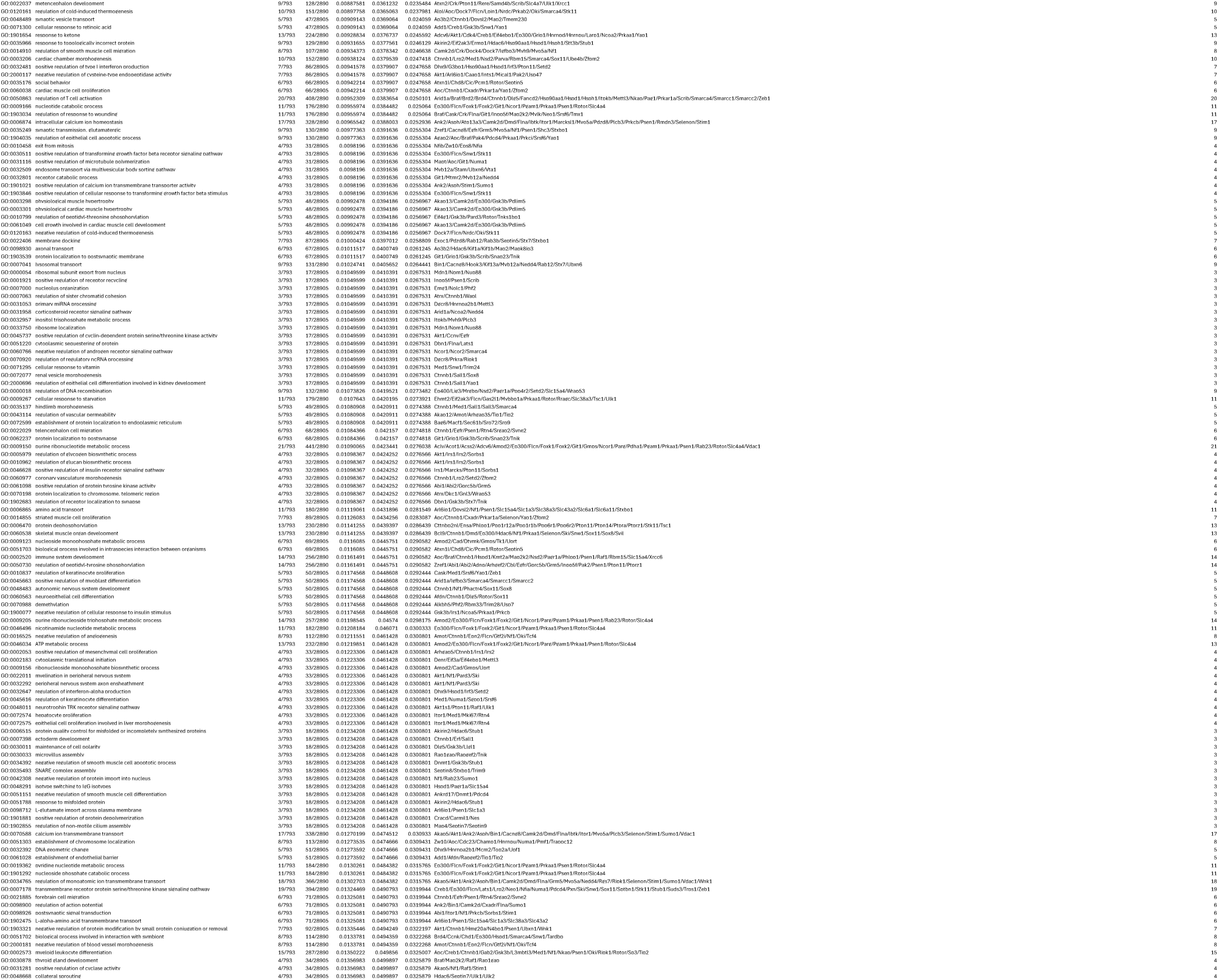
Spreadsheet_3_Tab_3\.

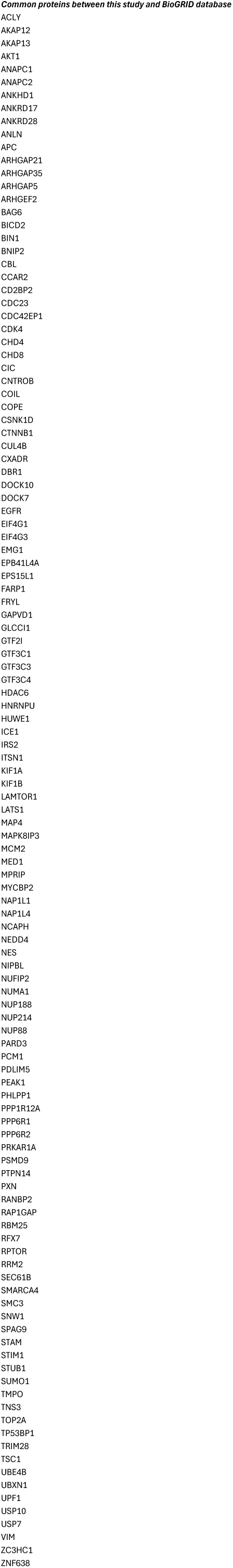
Supplemental_Spreadsheet_4.

Supplemental_Spreadsheet_5.

Primary antibodies used in this work
| Antigen (Species*) | Working dilution; use<sup>§</sup> | Origin |
| --- | --- | --- |
| **BrdU** | 1:500; IHC | Abcam ab6326 |
| **Cyclin D1 (M)** | 1:1000; IB | BD Biosciences 556470 |
| **Cyclin D2 (Rb)** | 1:1000; IB | Santa Cruz Biotechnology sc-593 |
| **DCX (Rb)** | 1:200; IHC | Abcam AB18723 |
| **FLAG-M2 (M)** | 1:1000; IB | Sigma-Aldrich F3165 |
| **GAPDH (M)** | 1:1000; IB | Sigma-Aldrich MAB374 |
| **GFAP (Ch)** | 1:500; IHC | Sigma-Aldrich AB5541 |
| **HUWE1 (Rb)** | 1:1000; IB / 5 µg; IP | Bethyl Laboratories A300-486A |
| **Ki67 (Rb)** | 1:200; IHC | Merk Millipore AB9260 |
| **Mash1 (Ascl1) (M)** | 1:100; IF / 1:1000; IB | BD Pharmingen 556604 |
| **MYC (M)** | 1:1000-1:5000; IB / 2.5 µg; IP / 1:500; IF | Sigma-Aldrich 05-724MG |
| **PLK1 (M)** | 1:1000; IB / 2.5 µg; IP | ThermoFisher 33-1700 |
| **PLK1 (M)** | 1:1000; IB | Abcam ab17057 |
| **Sox2 (Gt)** | 1:250; IHC | R&D Systems AF2018 |
| **VINCULIN (M)** | 1:10 000; IB | Sigma-Aldrich V9131 |
| **α-Tubulin (M)** | 1:10 000; IB | Sigma-Aldrich T9026 |
| **β-Actin (M)** | 1:40 000; IB | Sigma-Aldrich A2228 |
| **β-Catenin (Rb)** | 1:1000; IB | Cell Signalling Technology #9587 |

\*Rb, rabbit; M, mouse; Gt, goat; R, rat; Ch, chicken.
<sup>§</sup>IF, immunofluorescence; IHC, immunohistochemistry; IB, immunoblot; IP, immunoprecipitation

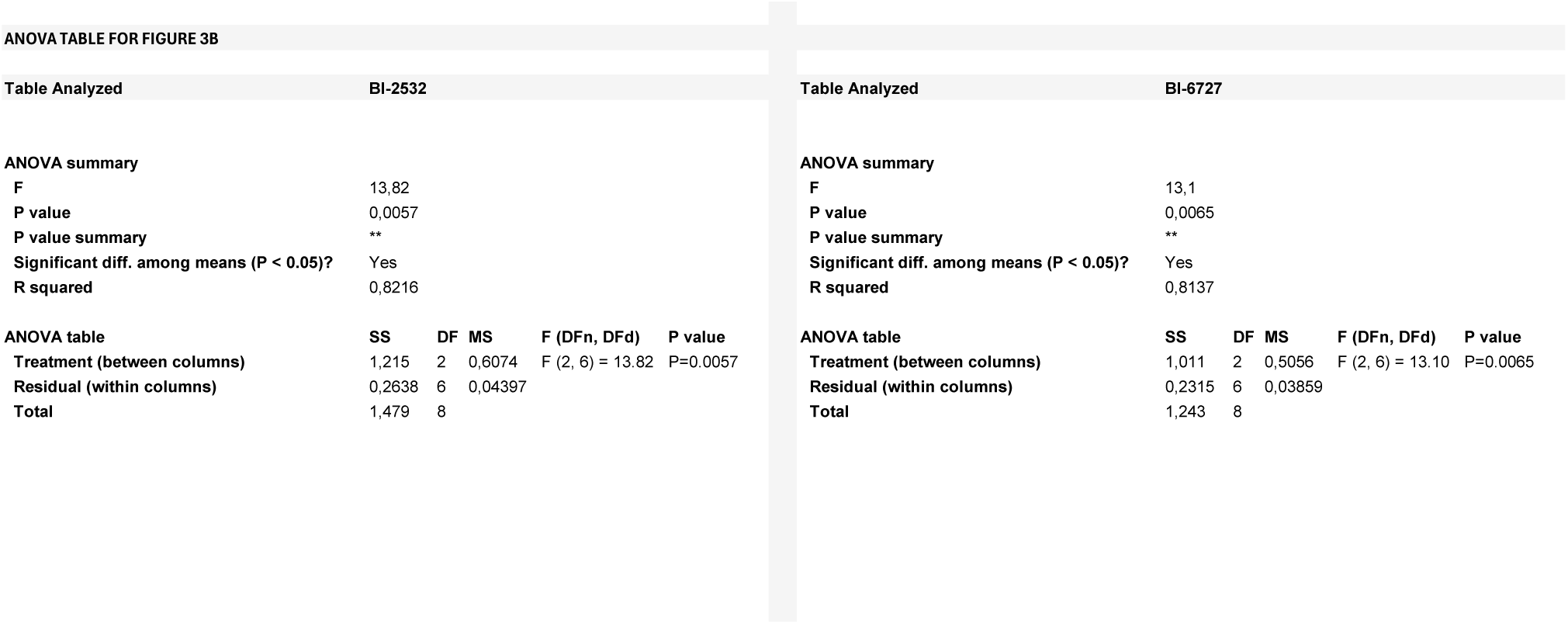
Supplemental_Spreadsheet_6_Tab_1.

Supplemental_Spreadsheet_6_Tab_2.

ANOVA TABLE FOR FIGURE 3G
| Table Analyzed | Effect of PLK1 mutants on Ascl1-myc |  |  |  |  |
| --- | --- | --- | --- | --- | --- |
| ANOVA summary |  |  |  |  |  |
| F | 23,58 |  |  |  |  |
| P value | <0,0001 |  |  |  |  |
| P value summary | **** |  |  |  |  |
| Significant diff. among means (P < 0.05)? | Yes |  |  |  |  |
| R squared | 0,7237 |  |  |  |  |
| ANOVA table | SS | DF | MS | F (DFn, DFd) | P value |
| Treatment (between columns) | 2,576 | 2 | 1,288 | F (2, 18) = 23.58 | P<0.0001 |
| Residual (within columns) | 0,9833 | 18 | 0,05463 |  |  |
| Total | 3,559 | 20 |  |  |  |

Supplemental_Spreadsheet_6_Tab_3.

ANOVA TABLE FOR FIGURE 3I
| Table Analyzed | Ascl1-myc normalised MFI |  |  |  |  |
| --- | --- | --- | --- | --- | --- |
| ANOVA summary |  |  |  |  |  |
| F | 7,181 |  |  |  |  |
| P value | 0,0051 |  |  |  |  |
| P value summary | ** |  |  |  |  |
| Significant diff. among means (P < 0.05)? | Yes |  |  |  |  |
| R squared | 0,6422 |  |  |  |  |
| ANOVA table |  |  |  |  |  |
|  | SS | DF | MS | F (DFn, DFd) | P value |
| Treatment (between columns) | 0,8572 | 3 | 0,2857 | F (3, 12) = 7,181 | P=0,0051 |
| Residual (within columns) | 0,4775 | 12 | 0,03979 |  |  |
| Total | 1,335 | 15 |  |  |  |

Supplemental_Spreadsheet_6_Tab_4.

ANOVA TABLE FOR FIGURE 3L
| Table Analyzed | Effect of PLK1 mutants and MG132 |  |  |  |  |
| --- | --- | --- | --- | --- | --- |
| Two-way ANOVA | Ordinary |  |  |  |  |
| Alpha | 0,05 |  |  |  |  |
| Source of Variation | % of total variation | P value | P value summary | Significant? |  |
| Interaction | 1,108 | 0,4216 | ns | No |  |
| PLK1 Mutant | 32,08 | <0.0001 | **** | Yes |  |
| Treatment (+MG132) | 48,13 | <0.0001 | **** | Yes |  |
| ANOVA table | SS | DF | MS | F (DFn, DFd) | P value |
| Interaction | 0,1005 | 2 | 0,05026 | F (2, 30) = 0.8890 | P=0.4216 |
| PLK1 Mutant | 2,911 | 2 | 1,456 | F (2, 30) = 25.75 | P<0.0001 |
| Treatment (+MG132) | 4,368 | 1 | 4,368 | F (1, 30) = 77.26 | P<0.0001 |
| Residual | 1,696 | 30 | 0,05653 |  |  |

